# Limitations of composability of cis-regulatory elements in messenger RNA

**DOI:** 10.1101/2021.08.12.455418

**Authors:** Samuel Haynes, Jamie Auxillos, Weronika Danecka, Abhishek Jain, Clemence Alibert, Edward Wallace

## Abstract

Genes are commonly abstracted into a coding sequence and cis-regulatory elements (CREs), such as promoters and terminators, and short sequence motifs within these regions. Modern cloning techniques allow easy assembly of synthetic genetic constructs from discrete cis-regulatory modules. However, it is unclear how much the contributions of CREs to gene expression depend on other CREs in the host gene. Using budding yeast, we probe the extent of composability, or independent effects, of distinct CREs. We confirm that the quantitative effect of a terminator on gene expression depends on both promoter and coding sequence. We then explore whether individual cisregulatory motifs within terminator regions display similar context dependence, using transcriptome-wide datasets of mRNA decay. To test the extent of composability, we construct reporter genes consisting of combinations of motifs within various terminator contexts, paired with different promoters. Our results show that the effect of a motif on RNA abundance depends both on its host terminator, and also on the associated promoter sequence. This emphasises the need for improved motif inference that includes both local and global context effects, which in turn could aid in the accurate use of CREs for the engineering of synthetic genetic constructs.

## INTRODUCTION

Our understanding of gene expression relies on the abstraction of a gene into discrete cis-regulatory elements (CRE), or parts, based on their most prominent function. The promoter coordinates the initiation of transcription. Transcription then progresses until it reaches the terminator, where cleavage and (in eukaryotes) polyadenylation of the nascent RNA transcript is coordinated. Extending the abstraction to the mature messenger RNA transcript, the coding sequence is flanked by untranslated regions (UTRs) at its 5’ and 3’ ends. The 5’UTR contributes to translation regulation and the 3’UTR contributes to transcript stability and localisation.

Abstracting specific characteristics to individual parts of a gene has facilitated the development of genetic toolkits that are based on the modular assembly of interchangeable parts. Furthermore, it is tempting to assume that a part with a function validated in a single context, e.g. in a specific gene and under certain experimental conditions, would have the same function in a different context. However, this assumption blurs two concepts of modularity: modular assembly, how easy it is to construct combinations of parts; and composability, how function and regulation can be decomposed into contributions from individual parts. The assembly of parts whose functions are not composable will result in constructs with unpredictable functions or regulatory behaviours.

Modular assembly is now routine: modern DNA synthesis and cloning techniques allow the construction of complex gene pathways through the combinatorial assembly of chosen parts (1–5). Several groups have constructed standardised libraries comprising promoter (including 5’UTR) and terminator (including 3’UTR) parts, which can be combined to achieve desired expression of synthesized proteins. Testing of standardised libraries tends to focus on one kind of part at a time, for example testing many promoters with the same coding sequence and terminator (2). Similarly, even massively parallel approaches to characterise smaller CREs usually explore a large library of promoter elements with a single terminator (6), or conversely a library of terminators with a single promoter (7). Because these experiments do not measure interactions between different parts, they rely on the untested assumption that those parts have more or less identical effects across different contexts.

Computational methods for the discovery and quantification of short CREs, such as sequence motifs recognised by regulatory proteins, also tend to rely on models of independent effects, that is, composability. Prominent methods for mapping CRE sequence-function relationships include predicting functional data with short sequence features, often using linear models (8–10); or, directly comparing the sequences of genes with similar characteristics to determine the presence of short consensus sequence motifs (11, 12) using motif discovery software (13, 14). Both of these approaches make the implicit approximation that the contribution of a short CRE is independent of context, so that the effect of combining motifs is composed of a linear sum (on the appropriate scale) of the individual CRE contributions. The approximation that short CREs act independently helps to find elements that have clear contributions, and to simplify a vast search space that would be made exponentially larger by accounting for CRE interactions. However, framing the search for CREs around independent contributions from short motifs overlooks multi-part motifs, interactions between motifs, and that motifs may be active only in specific contexts.

The importance of interactions among CREs has been clear since the discovery of cooperative DNA binding by transcription factors (15). For example, whether post-transcriptional regulatory CREs are functional or not depends on the RNA sequence context (16). One large-scale study of synthetic combinations of bacterial promoters and ribosomal binding sites showed that expression was largely composed of independent contributions; 64% of constructs had protein expression levels with less than a 2-fold deviation from the predictions of a linear model without interactions (17). However, 5% of the constructs had, on average, a 13-fold deviation. A recent study in budding yeast also found strong pairwise interactions between different promoter regions (18). Experiments with promoter-terminator swaps in budding yeast revealed 2-fold changes due to promoter-terminator interactions (3), even though these CREs are separated by hundreds of nucleotides of coding sequence.

Here, we provide further evidence for the importance of interactions between regulatory regions, and show the limitations of assuming that gene expression regulation can be decomposed into independent contributions from short CREs. First, we use modular cloning to construct 120 combinations of promoter (including 5’UTR), coding sequence, and terminator (including 3’UTR) modules, showing that the quantitative effect of terminators on protein production depends on both promoter and coding sequence. We then investigate the effect of promoter and terminator context on the regulatory behaviour of short CREs in 3’UTRs. In order to select suitable 3’UTR motifs, we adapt a method to computationally predict mRNA log2-half-life as a linear combination of codon usage and motif counts (9). We then quantify the effect on mRNA and protein levels of selected 3’UTR motifs when inserted in, or removed from, diverse reporter constructs. Our results demonstrate that the incorporation of candidate decay motifs indeed results in a decrease in mRNA abundance. However, the quantitative contribution from motifs depends on both the local 3’UTR context and the promoter context. These results highlight limits to the composability of cis-regulatory elements.

## MATERIALS AND METHODS

### Strains and media

*Saccharomyces cerevisiae* strain BY4741 (*MATa his3*Δ1 *leu2*ΔO *metl5*ΔO *ura3*Δ0) was used as the wild-type strain in this study, and the host for all yeast plasmid transformations. For all quantitative assays, plasmid-transformed strains were grown in synthetic complete medium without uracil (SC-Ura), containing 0.69% yeast nitrogen base without amino acids and with ammonium sulfate (Formedium, Norfolk, UK), 0.193% amino acid drop-out supplement mixture (Formedium, Norfolk, UK) and 2% glucose. To prepare BY4741 for transformation, we grew it in YPDA medium, containing 2% peptone, 1% yeast extract, 2% glucose and 0.004% adenine.

### Construction of chimeric reporter plasmids

All fluorescence reporter plasmids were constructed by Golden Gate assembly using the YeastFab system as described in (4). Promoters, coding sequences and terminators were either amplified from the yeast genome or synthesised by a commercial vendor (IDT) then cloned into a parts accepting plasmid (HcKan_P for promoters, HcKan_O for coding sequences and HcKan_T for terminators) by Golden Gate assembly using Bsa1-HFv2 (NEB). A detailed protocol for Golden Gate assembly is available at protocols.io, doi:10.17504/protocols.io.bkqrkvv6. Using these parts libraries, the promoters, coding sequences and terminators were assembled together into the transcription unit acceptor plasmid (POT1-ccdB) by Golden Gate assembly using Esp3I (NEB); these are low-copy centromeric plasmids with URA3 selection marker, as described in (4). Plasmid inserts were confirmed by Sanger sequencing (MRC PPU DNA Sequencing and Services, Dundee). DNA sequences used in this study are summarised in Supplementary Tables S4 and S5. Assembled plasmids were transformed into yeast BY4741 using lithium acetate transformation (20), and selected in SC-URA agar plates to isolate successful transformants.

The mCherry coding sequence is as used in (4), which in turn was amplified from the mCherry sequence in (6). The mTurquoise2 coding sequence is as used in (3).

### Fluorescence measurements: Plate reader analysis of strain growth and fluorescence

Yeast with plasmids were grown in a 96-well deep well plate (VWR) containing 100*μ*l of SC-Ura medium with 2% glucose and grown for ~12 hours at 30°C in a shaking incubator set at 250 rpm. The next day, the cultures were diluted to an OD of 0.2. For each sample, 3 technical replicates of 200μl were transferred to a 96-well black microtiter plate (Corning) and grown according to the protocol described in (21). The Tecan Infinity M200 series plate reader was set at the temperature of 29.9 (range of 29.4-30.4°C) with linear shaking (6 mm amplitude at 200-220 rpm). OD measurements were carried out at an absorbance wavelength of 595 nm with a measurement bandwidth of 9 nm with 15 reads. mCherry fluorescence measurements were carried out with an excitation wavelength at 585 nm and an emission wavelength of 620 nm (excitation bandwidth of 9 nm and emission bandwidth of 20 nm) with the gain set at 100. mTurquoise2 fluorescence measurements were carried out with an excitation wavelength at 434 nm and an emission wavelength of 474 nm (excitation bandwidth of 9 nm and emission bandwidth of 20 nm) with the gain set at 60.

Plate reader data were analysed using omniplate software(22). Omniplate accounts for autofluorescence and fits a model to the time series data using Gaussian processes to infer the time of maximum growth rate for each well. We minimised growth-dependent effects by using the fluorescence at maximum growth rate for all of our protein fluorescence experiments. Each fluorescence measurement was also normalised by OD to remove dependency on cell number, so every protein fluorescence measurement is recorded as fluorescence per OD at max growth rate. A detailed protocol for setting up and conducting the plate reader assay is available at protocols.io, doi:10.17504/protocols.io.bbicikaw.

### RNA measurements: Strain growth, RNA extraction, RT-qPCR, RNA-Seq and analysis

Yeast with plasmids were grown in a 24-well deep well plate plate (4titude) containing 1.5 ml of SC-Ura for ~20 hours at 30°C in a shaking incubator set at 250 rpm. The next day, the OD was diluted to a starting OD between 0.15-0.2 in a 12-column deep well reservoir plate (4titude) to a total volume of 7 ml. Diluted cultures were grown at 30°C in a shaking incubator set at 90 rpm to an OD of 0.5-0.7 then pelleted by centrifugation. Pelleted cells in the plate were stored in −80°C.

To extract RNA, we adapted a silica column DNA/RNA extraction protocol from Zymo Research (Irvine, California, USA). The pelleted cells were thawed and individually resuspended in 400 *μ*l of RNA binding buffer (Zymo), then transferred to 2 ml screw cap tubes containing zirconia beads, lysed using the Precellys Evolution homogeniser then pelleted by centrifugation for 1.5 minutes. The supernatant was transferred to a Zymo Spin IIICG column (Zymo) then centrifuged. The flow through was mixed with 1 volume of ethanol then transferred to a Zymo Spin IIC column (Zymo) and centrifuged. This flow through was discarded and 1 volume of DNA/RNA Prep buffer (Zymo) was added then centrifuged. The column was washed with 700 *μ*l of Zymo DNA/RNA Wash buffer (Zymo) then centrifuged. The column was washed a second time, but with 400 *μ*l of Zymo DNA/RNA Wash buffer (Zymo). The column was centrifuged once more to remove residual wash buffer in the column. Lastly, 30 *μ*l of nuclease free water was added to the column then eluted. All centrifugation steps in the RNA extraction protocol were carried out at 12,000g for 1 minute unless otherwise stated. A detailed protocol for yeast growth and RNA extraction is available at protocols.io, doi:10.17504/protocols.io.beetjben.

The quantity and quality of the RNA was measured using both a spectrophotometer (DS-11, DeNovix, Wilmington, Delaware, USA) and Fragment Analyser (Agilent). 4 *μ*g of RNA was treated with DNAse1 (Thermo) then inactivated using the RapidOut DNA removal kit (Thermo) according to the manufacturer’s protocol. 2.5 *μ*l of Random primer mix (NEB) was added to the mixture then separated into 2 PCR tubes (one for -RT and one for +RT) then denatured at 70°C followed by cooling on ice. Reverse transcription (RT) master mix was prepared, containing 2 *μ*l of First Strand synthesis buffer, 0.75 μl of 10mM dNTP mix, 1.5 *μ*l of nuclease free water, 0.25 μl of RNase inhibitor and 0.5 *μ*l of SuperScript IV Reverse Transcriptase (Invitrogen) per reaction. 5 *μ*l of the RT master mix was added to the denatured RNA then incubated at 25°C for 5 minutes then 55°C for 1 hour. The cDNA was diluted with 200 μl of nuclease free water.

Target cDNAs were measured by quantitative PCR with Brilliant III Ultra-Fast SYBR Green qPCR master mix (Agilent) using a Lightcycler 480 qPCR machine (Roche). We measured all +RT reactions in technical triplicate, and negative control -RT samples using one replicate. We used the manufacturer’s software to calculate quantification cycle (Cq) for each individual well using the fit points method, and exported both raw fluorescence and Cq data. All primer sets were thoroughly validated by serial dilution and by confirming amplicon size. Sequences are available in Supplementary Table S6.

RT-qPCR data was analysed using our tidyqpcr R package v0.3 (https://github.com/ewallace/tidyqpcr). For each biological replicate, ΔCq values were calculated by normalising the median mCherry Cq values by the median Cq values of the three reference genes (RPS3, PGK1 and URA3). For the constructs with motif insertions in terminators, ΔΔCq values were calculated by normalising mCherry ΔCq by that of control construct mod_NNN strains (with the corresponding promoter) for tRPS3 and tTSA1 consructs. For the constructs with motif deletions in terminators, ΔΔCq values were calculated by normalising mCherry ΔCq by that of the WT terminator (with the corresponding promoter) for tPIR1 constructs. Complete scripts for qPCR analysis, quality control, and figure generation are available online at https://github.com/DimmestP/chimera_project_manuscript/.

RNA-seq libraries were prepared using QuantSeq 3’ mRNA-Seq Library Prep Kit REV for Illumina (Lexogen Gmbh, Vienna Austria). See https://doi.org/10.1016/bs.mie.2021.03.020. 500 ng of RNA (not treated with DNaseI) was used as input and manufacturer’s protocol was followed without modifications. The number of amplification cycles was determined using PCR Addon Kit for Illumina (Lexogen). The quality of the library was measured using Fragment Analyzer NGS Fragment Kit (1-6000bp) (Agilent). Pooled libraries were then sequenced using NextSeq 500/550 (Illumina) with paired-end reads using Custom Sequencing Primer to obtain 3’-end reads.

5PSeq libraries were prepared as described in (23) with modifications to the reverse transcription step: anchored oligo(dT) was used instead of oligo(dT) to allow for sequencing of 3’-ends and random primers were not used. Library was sequenced using NextSeq system (Illumina) with paired-end reads.

RNA-Seq alignment and quality control was conducted using a pipeline available online at https://github.com/DimmestP/nextflow_paired_reads_pipeline, written in Nextflow (24). Quality control was conducted with FASTQC and MultiQC reports (25) and adapters were removed with Cutadapt (26). Alignment was conducted with HISAT2 (27), followed by processing with SAMtools (28) and BEDTools (29). The sacCer3 (R64-2) genome build was used for alignment and transcriptome annotation was originally taken from the Saccharomyces Genome Database (30). 5PSeq reads contain UMIs, which were used to deduplicate reads using UMI-tools (31); QuantSeq reads do not. Counts to genomic regions of interest were calculated using FeatureCounts (32). For 5PSeq data, 5’P ends were also analysed using fivepseq pipeline (33).

### Determining 3’UTR decay motifs

We initially selected 69 3’UTR motifs to investigate from three separate studies of cis-regulatory elements suspected to regulate mRNA decay (8,9,11). To select a short list of motifs to test for context dependence, we determined each motif’s contribution to a linear model predicting half-life. Following (9), we quantified the effect of motifs on transcript half-life using a linear model predicting half-life on the basis of codon usage, 3’UTR length, and 3’UTR motif frequency.

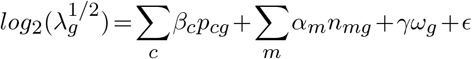

where 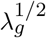 is the half-life of gene g, *β_c_* is the coefficient associated with codon c, *p_cg_* is the proportion of gene g’s coding sequence that corresponds to codon c, *γ* is the coefficient associated with 3’UTR length, *ω_g_* is the 3’UTR length of gene *g, α_m_* is the coefficient associated with motif *m, n_mg_* is number of occurrences of motif *m* in gene *g*’s 3’UTR, and *ϵ* is the noise term. To choose 3’UTR lengths and to assess which sequence to use for 3’UTR motif search, we used the median 3’UTR length estimates (precisely, the median length of clustered major transcript isoforms) reported from the TIF-seq analysis in (34).

We removed motifs that did not significantly contribute to half-life by using a greedy model selection algorithm that minimises the Akaike information criterion (AIC).

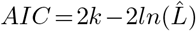

where *k* is the number of parameters in the model and *L* is the maximum value of the likelihood function (35). We implement this motif comparison using the R function step (36, 37), to iteratively add the motif which reduces the model’s AIC the most until the penalty for adding news terms overcomes the benefit of including a new motif. This procedure was run on each decay data set independently.

The variance explained by each predictor in the linear model trained on the (38) dataset is estimated in three ways. First, the variance explained by three individual linear models each containing one of the predicates is calculated. Then, we determine the reduction in variance explained when one of the predicates is removed from the full model. Finally, the increase in variance explained when each predicate is added to the model sequentially, starting with codon usage and ending with 3’UTR length, is reported.

We selected the specific versions of the HWNCATTWY and TGTAHMNTA motifs by running two addition linear models predicting half-life that inferred separate coefficients for each version of its consensus sequence. Coefficients were reported for the significant motif versions (Supplementary Table S7, S8). We chose instances with similar effect size, statistical significance, and number of occurrences in native transcripts. We chose TTTCATTTC for HWNCATTWY and for TGTAHMNTA chose TGTACAATA over TGTATATTA specifically to avoid the 5nt stretch also found in ATATTC.

### Design of modified 3’UTRs for testing the effects of mutated motifs

RPS3 was chosen as the first 3’UTR for inserting motifs into as it was the only terminator in the characterized library that did not contain any of the 69 original motifs of interest. The tRPS3 3’UTR-terminator was modified to incorporate three 9 nt insertion sites for motifs (M1, M2 and M3). The M1 was inserted 24 nt downstream of the stop codon, M2 was inserted 15 nt downstream of M1 and the final insert M3 was inserted 4 nt downstream of M2 (Figure 4A). These positions were selected based on key design criteria, including; minimal perturbations of RNA secondary structure as predicted by RNAfold (39), position of motifs in native 3’UTRs and position of other CREs important for transcriptional control (Supplementary Figure S3, Supplementary Table S9). A control tRPS3 3’UTR mod_NNN was designed to incorporate random bases in each insertion site. Further modified 3’UTR-terminator designs were designed to incorporate individual motifs of interest previously identified, within the insertion sites described (Figure 4A).

We chose an alternative 3’UTR for screening the effects of inserting motifs of interest by searching for characteristics similar to RPS3. To this end, median-length 3’UTRs were extracted from the (34) dataset filtered for the following criteria; 1) does not contain any of the original 69 motifs of interest, 2) < 300 nt in length, 3) from a highly expressed gene, 4) synthesisable as a gBlock by our manufacturer (IDT). The 3’UTR from TSA1 met these criteria.

Similar to the modified tRPS3 designs, in tTSA1 we designed three 9 nt motif insertion sites: M1 21 nt downstream of the stop codon, M2 20 nt downstream of M1, and M3 24 bp downstream of M2 (Figure 4A). The tTSA1 πiod_NNN construct contained random bases in the M1, M2 and M3 sites, with the motif insertions in other modified constructs as for tRPS3.

To design deletion constructs we selected a native 3’UTR that contained the motifs of interest. Again, median-length 3’UTRs were extracted from the Pelechano et al. (2013) dataset filtered for the following criteria; 1) contains at least 3 shortlisted motifs of interest, 2) a highly expressed gene, 3) synthesisable. The PIR1 terminator chosen for motif deletion contains one copy each of the ATATTC and UGUAHMNU motifs, and 3 copies of the HWNCAUUWY motifs (Figure 5A), although did not contain the putative stability motif GTATACCTA.

The mutation of motifs for their removal from the PIR1 3’UTR was carried out so that: 1) at least 50% of the motif sequence (specifically the motif consensus sequence) was mutated to a base that does not correspond to the consensus sequence, 2) GC content was minimally altered, 3) Mutations that resulted in a limited change in the predicted secondary structure and minimum free energy (MFE) according to RNAfold (39), see Supplementary Table S9).

The native and modified candidate 3’UTRs were screened for the presence of Esp3I and BsaI sites within the sequence. For incorporation into the YeastFab system, the sequence ‘agcgtgCGTCTCgTAGC’ was added to the 5’-end of the 3’UTR and the sequence ‘CCTCcGAGACGcagcac’ was added to the 3’-end of the 3’UTR. To check if sequences were synthesisable, 100 nt downstream of the native 3’UTR was added to the candidate and the sequence checked at the IDT gBlock entry form (https://eu.idtdna.com/site/order/gblockentry).

### Determining motif effect on abundance

A linear model predicting construct ΔCq’s using the presence or absence of the four selected motifs was trained on each promoter-terminator pairing separately. The model included a term to account for interactions between the UGUAMNUA and HWNCAUUWY motifs The linear model also included a term for batch effects, between the 2 experimental batches of 3 biological replicates for each set of constructs, because this improved the quality of model fit. The model was:

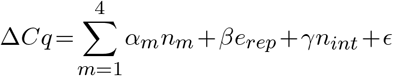

where *n_m_* is the copy number of motif m in the construct, *e_rep_* is which experimental batch the construct was part of and *n_int_* is the interaction term with value 1 if both UGUAMNUA and HWNCAUUWY motifs are present.

### Predicting changes in transcript abundance from changes in half-life

A simple kinetic model of the production and decay of transcripts was used:

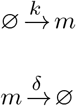

The steady state solution for the average number of transcripts, 〈*m*〉, is

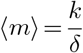

where k is the rate of transcription, which can include multiple active states, and *δ* is the rate of decay for the transcript (40).

Now consider a control transcript *m*_0_, and a similar transcript with an altered terminator *m_a_*. Assuming the alterations to the host gene’s terminator have a minimal impact on the transcription rate, the above equation says that the ratio of predicted abundance 〈*m_a_*〉 to the control transcript abundance, 〈*m*_0_〉, is the same as the ratio of their half-lives:

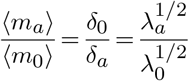

This gives a linear effect on the log-scale abundance

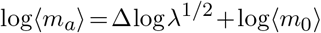

and because PCR quantification cycle Cq is proportional to log2(*m*), this directly leads to a linear effect on Cq.

All analysis made extensive use of the tidyverse and ggplot2 (41, 42).

## RESULTS

### Terminator effects on gene expression depend on cis-regulatory context

To investigate the context dependence of cis-regulatory regions, we created a library of 120 constructs (Figure 1A), containing all combinations of 6 promoters (including 5’UTR), 2 coding sequences, and 10 terminators (including 3’UTR). We selected promoters and terminators from native yeast genes spanning a variety of different expression patterns and functions (Table 1). To choose specific sequence lengths for the terminators of our constructs, we referred to published measurements of median 3’UTR length (34) because sequences that are necessary and sufficient for efficient transcriptional termination are found upstream of the termination site (43). For terminators with measured length under 200nt, we used the standardised parts length of 200nt from the YeastFab library. The 2 coding sequences (CDS) expressed mCherry or mTurquoise2 fluorescent proteins, which are bright fluorophores with only 30% amino acid identity (44, 45).

**Figure 1.**
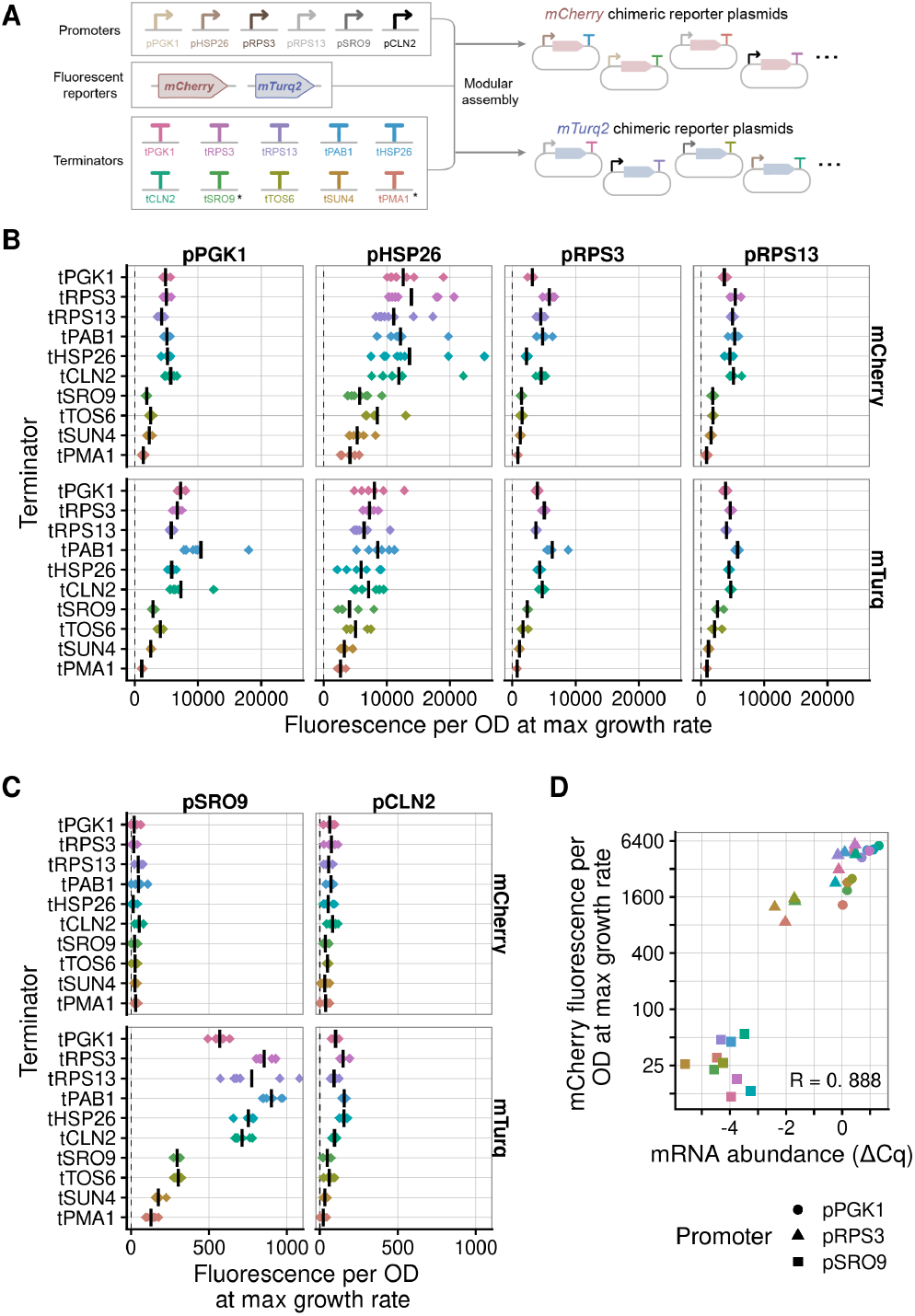
Both terminator and promoter contribute to gene expression. (**A**) Design of chimeric reporter constructs with all combinations of 6 promoters, 2 fluorescent proteins, and 10 terminators, on a centromeric plasmid. (**B**) Protein abundance estimated by mCherry and mTurquoise2 (mTurq) fluorescence for 10 terminators paired with 4 high expressing promoters. Fluorescence and OD were measured in cultures grown in a plate reader, and reported at the time of each sample’s maximum growth rate (see methods). Each diamond represents a biological replicate, averaged over 3 technical replicates. The vertical line is the mean of all 6 biological replicates. (**C**) Same as panel B, but for 2 low expressing promoters. Negative fluorescence values arising from instrument noise dominating measurements of constructs with negligible fluorescence are automatically set to 0. (**D**) mCherry fluorescence correlates with RT-qPCR mRNA abundance for 3 promoters paired with all 10 terminators. Note that both axes use a *log*_2_ scale.

**Table 1.** Summary of the terminator library. The common gene name from which the terminator is extracted from is included alongside its systematic name. The median 3’UTR reported by (34) is a median over the lengths of each distinct isoform they detect. The usage of each motif is signified by S, for promoter-terminator swaps, or M, for motif insertion or deletion. A short summary of the protein function is included.

| Gene Name | Systematic Name | Median 3'UTR Length | Construct Terminator Length | Usage | Function |
| --- | --- | --- | --- | --- | --- |
| PGK1 | YCR012W | 158 | 189 | S | Glycolysis |
| RPS3 | YNL178W | 86 | 200 | M&S | Ribosomal |
| RPS13 | YDR064W | 92 | 200 | S | Ribosomal |
| PAB1 | YER165W | 150 | 200 | S | RNA Binding |
| HSP26 | YBR072W | 164 | 200 | S | Heat Shock |
| CLN2 | YPL256C | 203 | 200 | S | Cell Cyclin |
| SRO9 | YCL037C | 543 | 545 | S | RNA Binding |
| TOS6 | YNL300W | 256 | 256 | S | Cell Wall |
| SUN4 | YNL066W | 198 | 198 | S | Cell Wall |
| PMA1 | YGL008C | 421 | 421 | S | Transmembrane ATPase |
| TSA1 | YML028W | 112 | 219 | M | Redox Homeostasis |
| PIR1 | YKL164C | 235 | 358 | M | Cell Wall |

Measuring fluorescence with a plate reader showed that, as expected, promoter choice dominated overall protein output. We observed 100-fold changes in fluorescence between the 4 highest expressing promoters (Figure 1B) and the 2 lowest expressing promoters (Figure 1C) across both coding sequences. Expression from the stress-induced pHSP26 was notably more variable than from pPGK1 and pRPS’s, across 6 biological replicates, when combined with both coding sequences and a variety of terminators. We also confirmed that most differences in protein outputs are accounted for by changes in mRNA abundance by checking a subset of mCherry constructs using RT-qPCR (*R*=0.888, Figure 1D).

Terminators also affect protein output with 5-fold changes in fluorescence seen within the same promoter-CDS sets, relative to the tPGK1 terminator of each group (Figure 2A). We focus on constructs with high-expression promoters due to the poor signal to noise ratio at low expression levels. The interaction of coding sequence and terminator is seen most clearly for tPAB1. tPAB1 is consistently the most highly expressed terminator in mTurquoise2 constructs, but is more variable in mCherry constructs. Meanwhile, tPGK1 highlights the interactions of promoter and terminator. tPGK1 is one of the most highly expressing terminators when paired with pPGK1 and pHSP26, but is up to 40% lower in expression when paired with pRPS3 or pRPS13. Overall, our results show that the contributions of terminators (including 3’UTRs) to gene expression depend on other parts within the gene.

**Figure 2.**
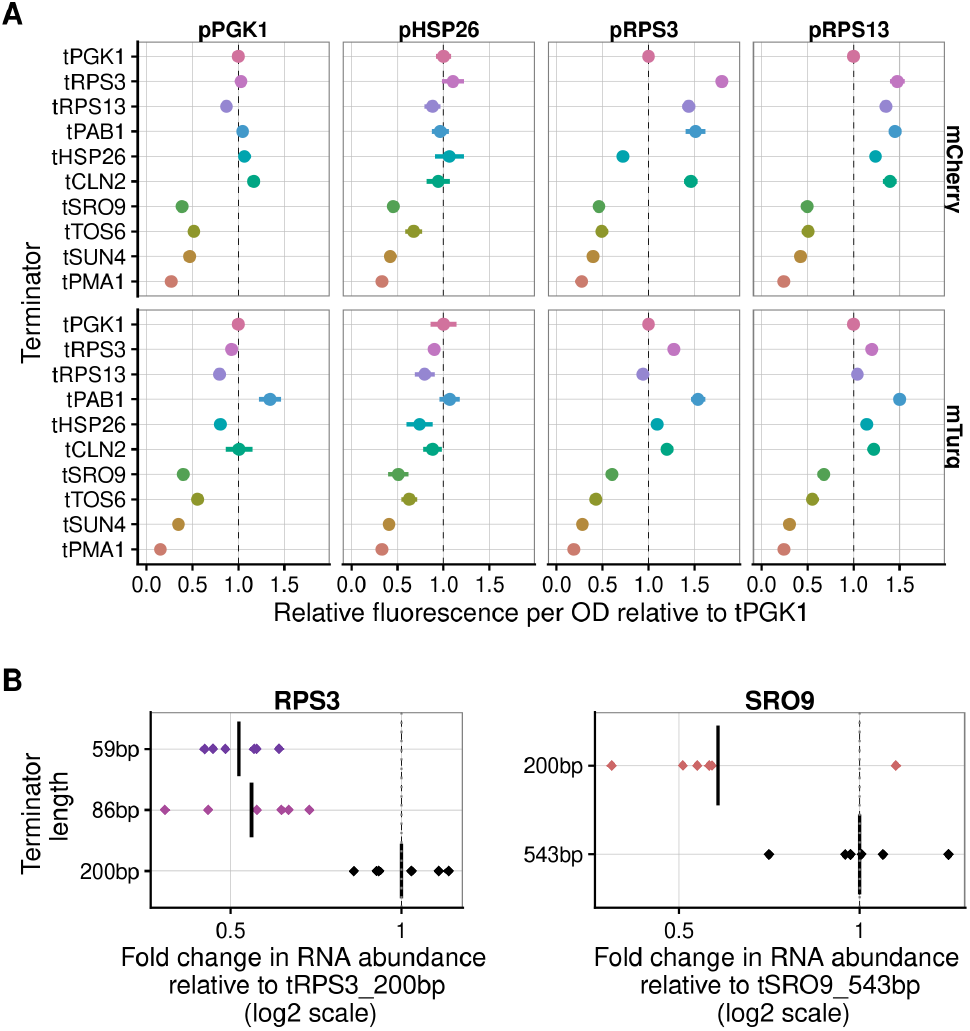
Terminator contributions to gene expression are promoter, coding sequence and length dependent. (**A**) Relative protein abundance from each terminator, normalized to a reference terminator tPGK1 for each matched promoter. The plot shows statistical summaries (mean and standard error) of 6 replicates for high-expression promoter data shown in Figure 1A. (**B**) RT-qPCR mRNA results targeting the mCherry ORF for pRPS3-mCherry-tRPS3 and pSRO9-mCherry-tSRO9 constructs with differing terminator lengths.

We further investigated the effects of terminators on mRNA levels by comparing constructs with extended vs truncated terminators. It is known that disrupting the transcription termination signal lowers expression (7, 43). However, standardised parts libraries often assume a fixed terminator length for all genes, which is likely to omit the termination signal for genes with longer terminators. In the case of the YeastFab parts library (2) the fixed terminator length is 200bp. We compared a gene with a median 3’UTR length less than 200bp, RPS3 to a gene with a median 3’UTR length greater than 200bp, SRO9. The median length of the native RPS3 3’UTR is 86nt (34); truncating the terminator to 86bp or 59bp reduces protein output by almost 2-fold (Figure 2B). The median length of the native SRO9 3’UTR is 543nt (34); extending the terminator length to 543bp increases protein output by almost 2-fold (Figure 2B). This validates our assay’s ability to detect known regulatory signals affecting transcription termination, while highlighting the importance of using well-informed annotations to construct parts libraries for synthetic biology. Note that we used the longer 543bp SRO9 terminator, and a similarly extended 421bp PMA1 terminator, for the main set of constructs (Figure 1).

### Candidate cis-regulatory elements contribute to transcript decay rates

Next, we investigated how the regulatory effects of CREs contained within terminator regions depend on their context. First, 69 suitable CREs to test for context dependence were found through a literature search. All were suspected sequence motifs for mRNA binding proteins, several directly associated with proteins involved in mRNA degradation (8, 9, 11). Any motifs that were found in fewer than 6 gene 3’UTRs, as annotated by (34), were removed.

We quantified the regulatory effects of the remaining 38 candidate motifs by applying a linear model predicting half-life to 2 recent transcriptome-wide analyses of mRNA decay that used metabolic labeling (38, 46). These datasets are loosely correlated in their half-life measurements across 4188 genes reported in both datasets, *R*=0.63 (Figure 3A). However, (38) estimated substantially smaller half-lives. (38) also had greater coverage of genes in the yeast genome, 5529 vs 4304, and used multiple time points to determine halflives. Following (9), we constructed a linear model to predict a transcript’s half-life using the counts of motifs in its 3’UTR, the length of the 3’UTR, and the relative codon usage in each transcript’s coding sequence (see Material and Methods). The linear model performed similarly on both datasets by explaining 44% and 41% of the variability in half-lives for the (38) and (46) datasets respectively (Figure 3C). This predictive power is comparable to the squared correlation between the datasets (*R*^2^ = 0.40). Motifs that did not significantly contribute to the model were automatically filtered out using a greedy algorithm maximising the Akaike information criterion (AIC) during both training stages. Approximately 1.7% of the variance is explained by 7 significant motifs, with 42.0% explained by codon usage (Supplementary Table S1), consistent with previous analyses (9, 47). The top 7 most significant motifs from the (38) data showed similar regulatory behaviour when tested on their own in the (46) data, except for TGTAAATA which was stabilising in one dataset and destabilising in another, as we later discuss (Figure 3B).

**Figure 3.**
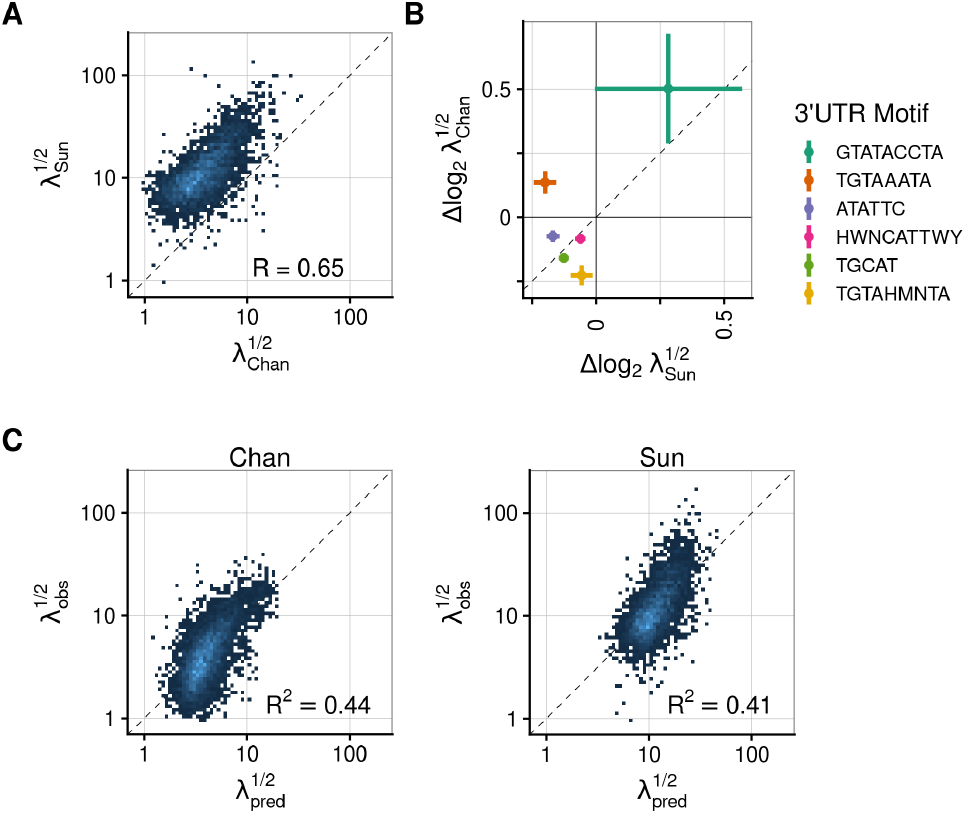
A linear model of transcript half-life quantifies the effect of candidate terminator motifs on half-life. (**A**) Correlation between the 2 transcript half-lives (λ), in minutes, reported in the (38) and (46) datasets. (**B**) Predicted contributions to log2 half-life for chosen motifs in the (38) and (46) datasets. (**C**) Predicted vs actual transcript half-lives calculated by a linear model of codon and motif usage trained on the (38) and (46) datasets.

We selected 4 motifs for exploring context dependence: TGTAHMNTA, GTATACCTA, HWNCATTWY, and ATATTC (Table 2). TGTAHMNTA and GTATACCTA were chosen as they had the largest coefficients amongst significant decay and stability motifs, respectively. HWNCATTWY was chosen due to its statistically significant effect in both datasets and, as it co-occurs with TGTAHMNTA in 68 native 3’UTRs, because it could be used for testing motif interactions. The final selected motif was ATATTC, as it is a statistically significant decay motif in both datasets, and it has been previously shown to lower mRNA abundance when inserted in reporter constructs (9). Functionally, TGTAHMNTA is the binding motif for Puf4p, and HWNCATTWY is associated with Khd1p/Hek2p-bound transcripts (11). However, it is not known how ATATTC and GTATACCTA affect mRNA decay.

**Table 2.**
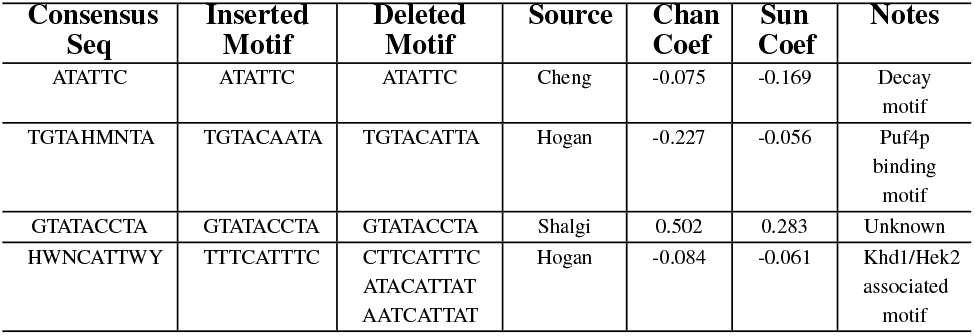
Summary of shortlisted motif characteristics. The first 3 columns hold the consensus sequence for each motif and the exact versions deleted from or inserted into the host terminators. Then, we report the paper from which the motif was selected from; (11), (8) or (9). Next, each motif’s coefficient given by the linear model predicting either the (38) or the (46) half-life datasets are included. Finally, the table includes notes on motif functions.

### Quantification of differential expression due to motif insertion or mutagenesis in multiple 3’UTRs

To quantify the effects and composability of selected motifs in different contexts, we designed a further set of reporter constructs (Figure 4A). We first chose the ribosomal protein terminator tRPS3, as it was the only terminator in our initial library that did not contain any of the selected motifs. We selected thioredoxin peroxidase tTSA1 as the second host terminator because it also lacks selected motifs and has similar length to tRPS3. In each host terminator we chose 3 motif insertion sites, selecting for: minimum impact on transcript secondary structure, avoiding known transcription termination elements, and matching the positions of motifs in native genes. Having 3 insertion sites enabled us to quantify combinations of motifs, including duplicates of weaker motifs to increase the likelihood of detecting a clear effect on gene expression. We chose TGTACAATA and TTTCATTTC sequences as explicit versions of the TGTAHMNTA and HWNCATTWY consensus motifs respectively, and checked that these explicit versions have similar predicted effects on half-life transcriptome-wide (Supplementary Tables S7, S8). Altogether, 7 variant terminators were designed for these 2 host terminators: the wildtype terminator, a control to test the insertion sites with randomly generated sequences, 4 testing the effects of inserting each motif individually and a final variant to test interactions between the TGTAHMNTA and HWNCATTWY motifs. We created a construct library by pairing each terminator with three different promoters; its native promoter pairing (pRPS3 or pTSA1), the high-expression promoter pPGK1, and the low-expression promoter pSRO9.

**Figure 4.**
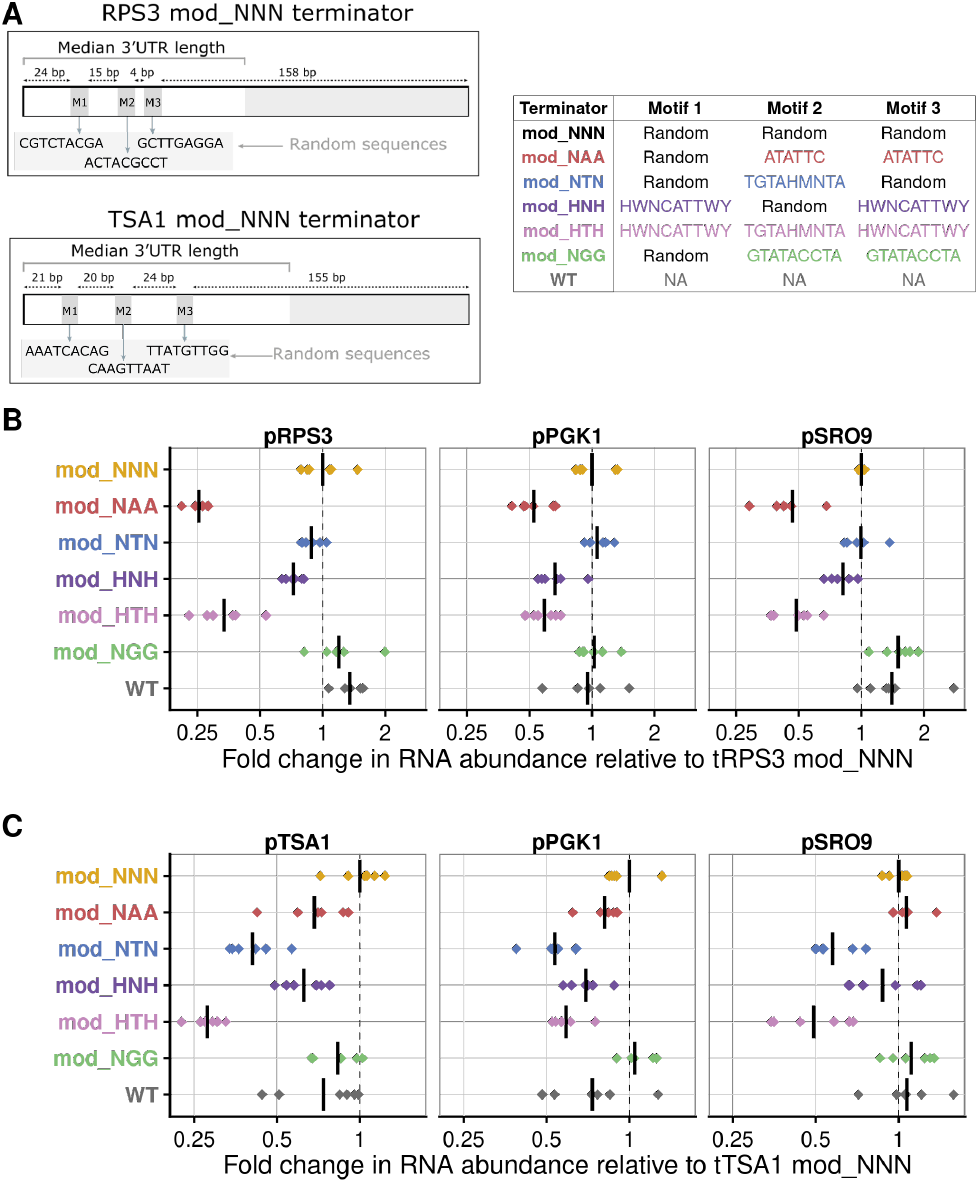
Motifs inserted into RPS3 and TSA1 host terminators change transcript abundance in RT-qPCR measurements. (**A**) Design of motif insertion sites in native RPS3 and TSA1 terminators, highlighting random insertion used as a negative control. (**B**) Fold changes in transcript abundance for tRPS3 constructs paired with 3 promoters: pRPS3, pPGK1 and pSRO9. (**C**) Fold changes in transcript abundance for tTSA1 constructs paired with three promoters: pTSA1, pPGK1 and pSRO9. Each diamond represents a biological replicate, averaged over 3 technical replicates. The vertical line represents the mean of all 6 biological replicates. Fold changes are relative to the abundance of the mod_NNN construct, i.e. 2^ΔΔ*Cq*^ (see methods).

Motifs predicted to affect half-life affect mRNA abundance when inserted into tRPS3 in reporter constructs (Figure 4B). We measured mRNA abundance by RT-qPCR across 6 biological replicates, each quantified in 3 technical replicates and normalised by the *ΔCq* method against values from 3 reference mRNAs (see methods). Insertion of 2 copies of ATATTC (mod_NAA) generally lowers the mRNA abundance, as much as 4-fold when paired with the pRPS3 promoter. Insertion of either TGTAHMNTA (mod_NTN), or 2 copies of HWNCATTWY (mod_HNH), tends to decrease mRNA abundance, and their combined insertion (mod_HTH) tends to decrease mRNA abundance even further. The putative stability motif GTATACCTA (mod_NGG) does not consistently or strongly affect mRNA abundance. However, comparison of the WT and control (mod_NNN) terminators does show that the creation of the insertion sites alone has an effect on mRNA levels.

Inserting the same motifs into our second host terminator gives qualitatively similar results (Figure 4C). Decay motifs generally lead to decay, although ATATTC (mod_NAA) has a weaker effect in tTSA1 than in tRPS3, and TGTAHMNTA (mod_NTN) has a stronger effect in tTSA1 than tRPS3. The putative stability motif GTATACCTA (mod_NGG) again has little effect.

We next quantified the effects of removing decay motifs from a native yeast terminator. We selected the cell wall protein PIR1 as our host terminator as it is only 258 bp ((34)) and a de-novo-synthesizable terminator that contains the ATATTC, TGTAHMNTA, and HWNCATTWY motifs. We designed 8 terminators in which the motif occurrences in tPIR1 were replaced by scrambled sequences (Figure 5A). We found that the removal of almost any decay motif from tPIR1 results in an increase in mRNA levels (Figure 5B).

**Figure 5.**
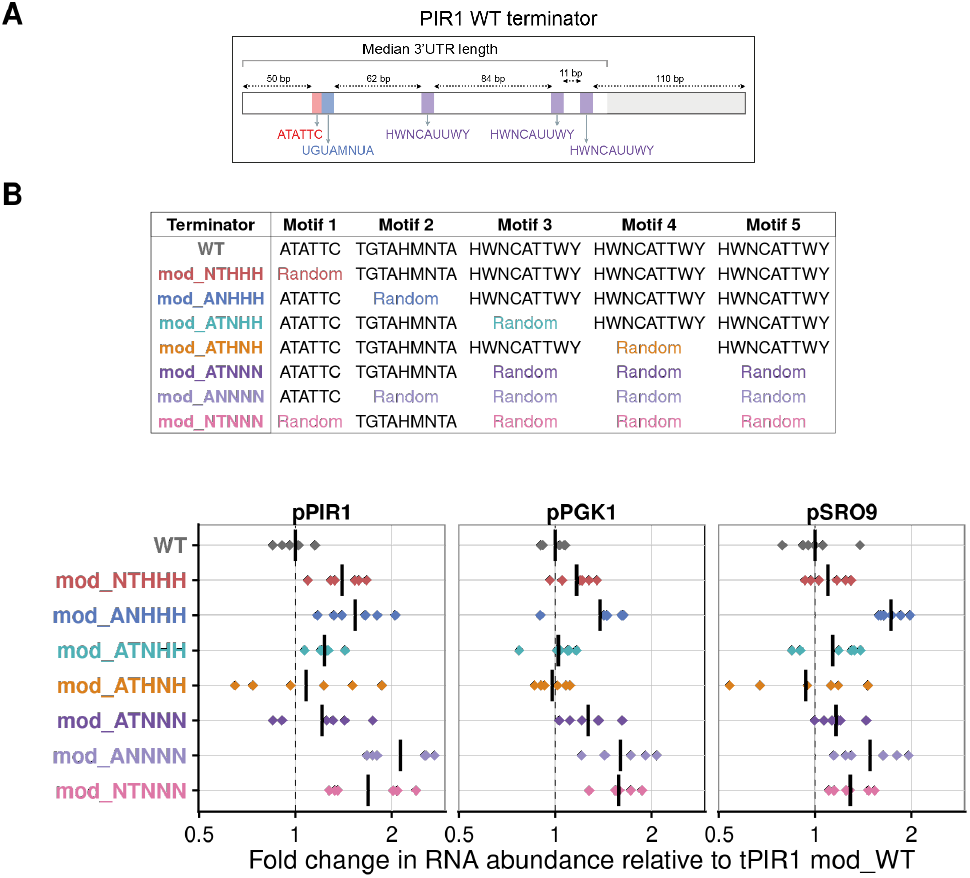
Motifs removed from PIR1 host terminators change transcript abundance in RT-qPCR measurements. (**A**) Design of PIR1 constructs with combinations of motifs replaced by random nucleotide sequences. (**B**) Fold changes in transcript abundance for tPIR1 constructs paired with 3 promoters: pPIR1, pPGK1 and pSRO9. Each diamond represents a biological replicate, averaged over 3 technical replicates. The vertical line represents the mean of all 6 biological replicates. Fold changes are relative to the abundance of the WT construct, i.e. 2^ΔΔ*Cq*^ (see methods).

We confirmed that motif-dependent changes in mRNA abundance are reflected in protein abundance by measuring the fluorescence from a subset of reporter constructs with native promoter-terminator pairings (Supplementary Figure S2). The high correlation (R=0.96,0.68,0.96 for tRPS3, tTSA1, tPIR1 constructs respectively) demonstrates that these combinations of decay motifs that change mRNA abundance also change the protein output, as expected.

Comparison of mRNA abundance across all constructs (Figure 4B, 4C, 5B) shows motif contributions change in magnitude but not direction depending on the context of the rest of the construct. The insertion of almost any decay motif into tTSA1 or tRPS3 results in a decrease in mRNA abundance, and removal of these from from tPIR1 results in an increase in mRNA abundance. However, the quantitative effects vary depending both on the immediate motif context in the host terminator and on the more distant context given by the promoter.

### Motif effects on gene expression depend both on terminator context and promoter pairing

We compared the effects of cis-regulatory motifs on mRNA abundance to predicted effects inferred from the transcriptome-wide measurements of half-life. First, we trained a linear model using the RT-qPCR results to estimate the change in log2 mRNA abundance (i.e. Δ*Cq*) due to the presence of a motif in each promoter and terminator combination. Using a simple model of transcript production and decay, we can argue that changes in mRNA abundance are directly proportional to changes in mRNA half-life (see methods). Therefore, we compared the motif’s estimated change in log2 mRNA abundance to that predicted due to changes in log2 half-life, estimated from our transcriptome-wide analysis of the (38) dataset. The effect of motifs in reporter constructs is correlated with the predictive model, but the strength of the correlation depends on context (Figure 6A). Constructs with inserted motifs (i.e. tRPS3 and tTSA1 constructs) had a lower correlation with predicted effects than constructs with deleted motifs (i.e. tPIR1 constructs). Interestingly, motif effects on mRNA abundance appear to be greater than that predicted from their effect on half-life when their host terminator is paired with its native promoter.

**Figure 6.**
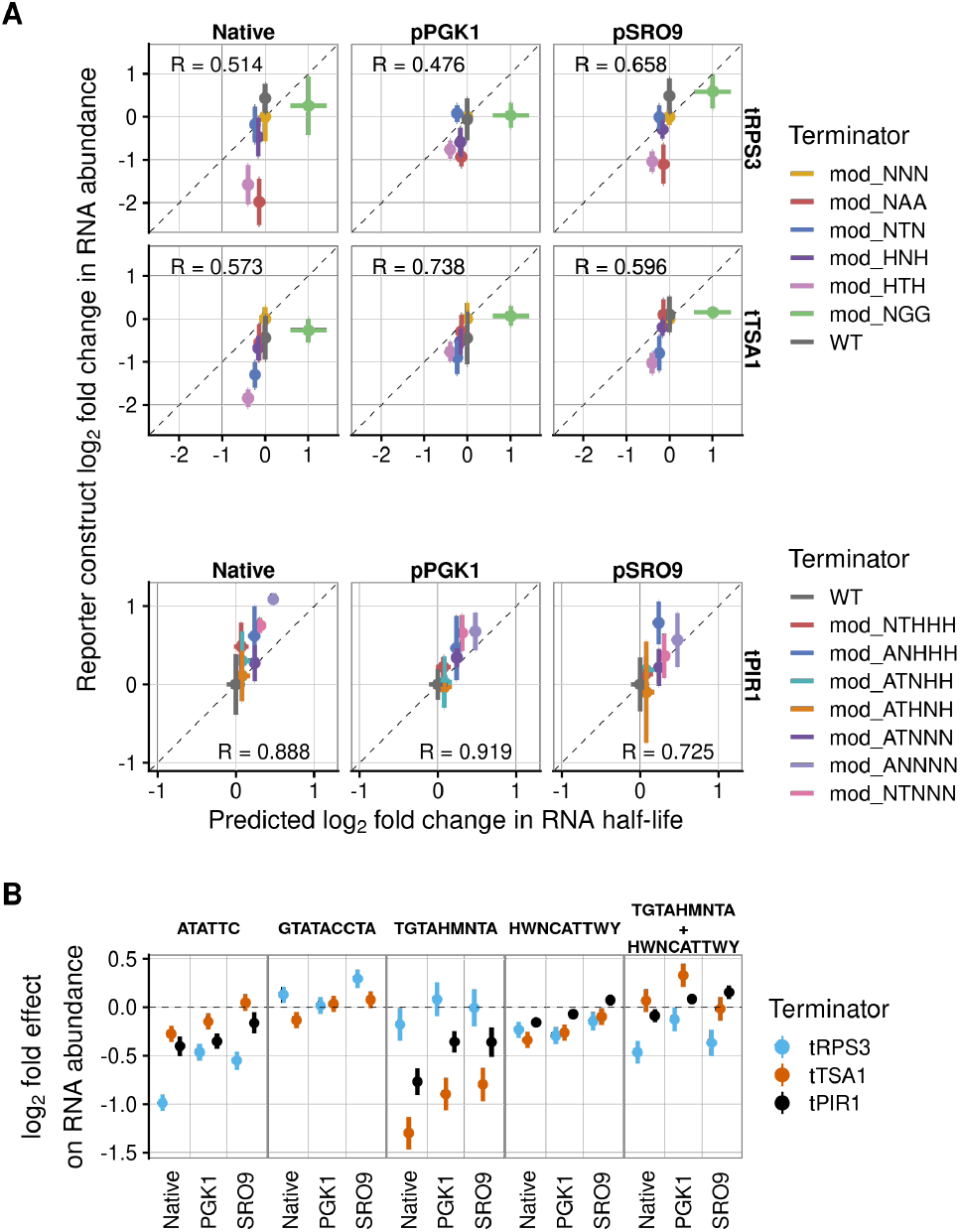
Promoter and terminator context alter the regulatory effect of motifs. (**A**) Predicted change in transcript abundance inferred from transcriptome-wide motif contributions to half-life, compared to RT-qPCR measurements of reporter transcript abundance. The y-axis shows statistical summaries (mean and standard error) of 6 replicates for data shown in figures 4 and 5. The fold change is relative to the mod_NNN construct in each promoter-terminator pairing for the tRPS3 and tTSA1 sets, and relative to the WT constructs for tPIR1 sets. Native promoter panels show the promoter paired with the terminator from the same set, e.g. pRPS3 with tRPS3. (**B**) Motif contributions to fold changes in mRNA abundance for reporter constructs with different promoter and terminator contexts. This is calculated by a linear model with a coefficient for the effect of each motif in each set, applied to *ΔCq* against 3 reference genes. The last column shows the interaction term between TGTAHMNTA and HWNCATTWY.

We next directly compared the estimated coefficients for each motif’s effect on mRNA abundance across promoter-terminator pairing (Figure 6B). The effect of a motif depends on terminator context. For example, ATATTCA reduces mRNA abundance substantially more when inserted in tRPS3 than in tTSA1. Meanwhile, TGTAHMNTA significantly reduces mRNA abundance when inserted in tTSA1, but not tRPS3, whichever promoter is chosen. Promoter choice also influences the magnitude of a motif’s contribution to mRNA levels. For the ATATTC, TGTAHMNTA and HWNCAUUWY motifs the greatest reduction in mRNA abundance occurred when native promoter-terminator pairings are measured. This is true for all 3 decay motifs across all 3 host terminators, except for HWNCAUUWY in pRPS3-tRPS3 constructs.

Regulatory interactions between different motifs also change depending on host terminator and promoter context. We included an interaction term that quantifies how the effect of including both HWNCAUUWY and TGTAHMNTA together differs from the sum of the effects of including these motifs individually. The combination of TGTAHMNTA and HWNCAUUWY in tRPS3 has no additional effect beyond a simple sum of their individual effects when paired with pPGK1. However, when tRPS3 is paired with pRPS3 or pSRO9, the combination has a greater effect than expected from the sum of each motif’s individual effect. The combination of TGTAHMNTA and HWNCAUUWY in tTSA1 has no additional effect, except when paired with pPGK1, where it has a lesser effect than expected. Finally, the combination of TGTAHMNTA and HWNCAUUWY in tPIR1 always has an additional effect across the three promoters. The combination has a greater effect than expected when paired with pPIR1 but a lesser effect when paired with pPGK1 or pTSA1.

### Inserting motifs into terminators shifts poly(A) site usage downstream

We next mapped poly(A) site usage in a subset of reporter constructs, for 2 reasons. First, changes in poly(A) site usage might mean that motifs placed in our reporters were unintentionally absent from the mature mRNA. Second, we wanted to know if motif effects on mRNA abundance might be due to changes in the poly(A) site usage. We chose three constructs with large effect sizes in the qPCR results: mod_NAA, mod_HTH and mod_NTN, together with WT and mod_NNN controls, within three promoter-terminator contexts: pRPS3-tRPS3, pPGK1-tRPS3 and pTSA1-tTSA1. For these constructs we performed paired-ends sequencing of 3’ mRNA-Seq libraries following QuantSeq protocol ((48) and see methods). Read 1 allows precise inference of poly(A) site position while read 2 generally overlaps the CDS and allows distinguishing terminators in native loci from reporter constructs. We mapped the reads to genome sequences extended by the relevant reporter plasmid sequence.

We detected 1000s of reads on each reporter construct, which is enough to quantify expression confidently as well as to assign poly(A) sites. We checked that counts of all other RNAs are highly correlated between samples, giving us confidence that changes in construct detection are meaningful (Supplementary Figure S11). Transcript abundance, relative to mod_NNN, for most constructs correlates strongly with qPCR results (Figure 7A). However, some constructs were detected as more abundant by RNA-seq for reasons that are unclear.

**Figure 7.**
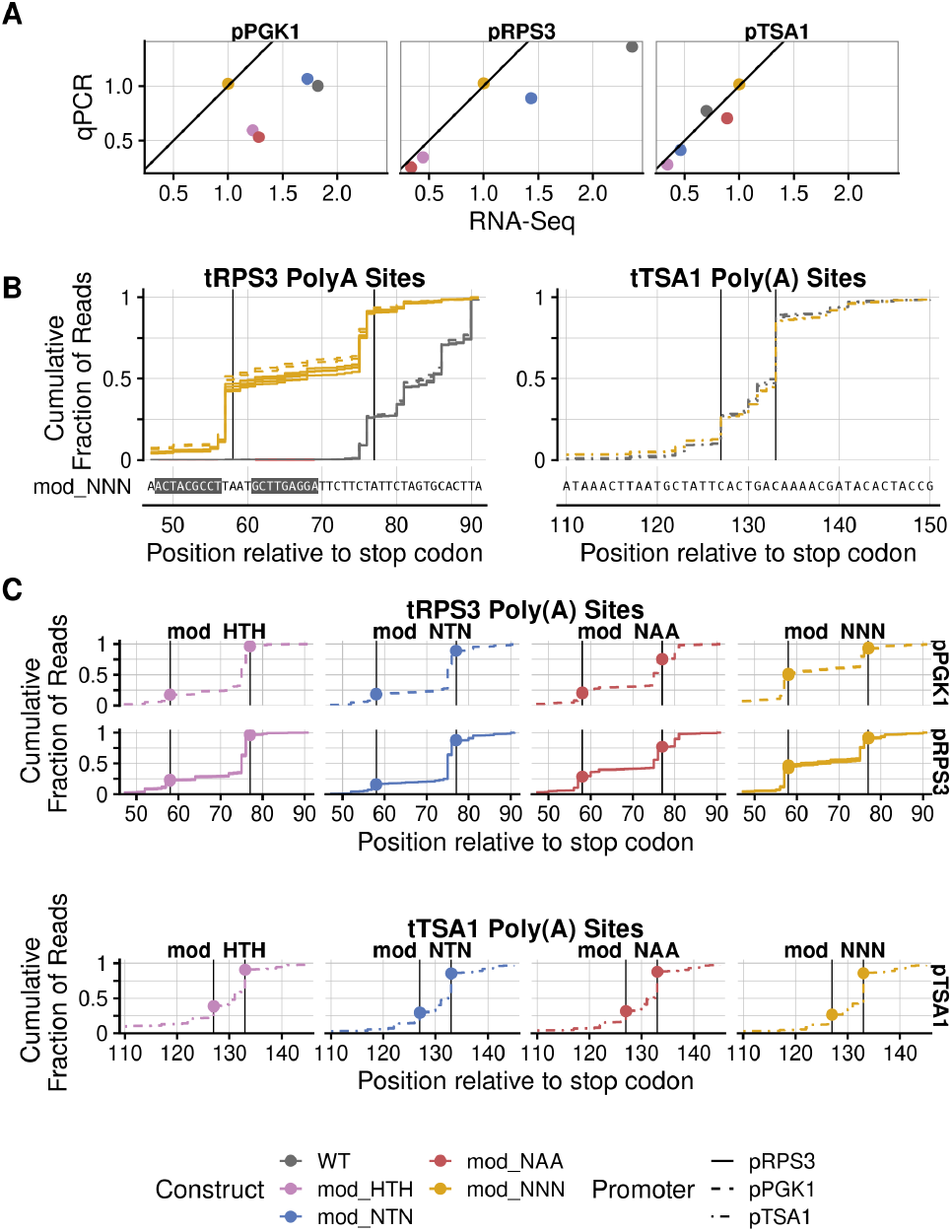
Inserting motifs into RPS3 and TSA3 terminators changes 3’UTR length. A subset of constructs were chosen to investigate changes in poly(A) site usage: WT, mod_NNN, mod_NTN, mod_NAA, and mod_HTH; within three promoter-terminator contexts: pRPS3-tRPS3, pPGK1-tRPS3, and pTSA1-tTSA1. (**A**) Comparison of construct transcript abundance as independently measured by qPCR and RNA-Seq assays. Transcript abundance was normalised to the median abundance of plasmid URA3, genomic PGK1, and RPS3 or TSA1 transcripts for each construct. Fold change is relative to the mod_NNN construct in each promoter-terminator context. The black diagonal line represents the expected values if RNAseq and qPCR results correlated perfectly. (**B**) Cumulative count of reads mapped downstream of WT (grey) and mod_NNN (golden) construct stop codons as fraction of total reads mapped to the constructs terminator. (**Left**) shows the poly(A) sites for the pRPS3-tRPS3 and pPGK1-tRPS3 promoter-terminator constructs. (**Right**) shows the same for the pTSA1-tTSA1 constructs. WT reads have been shifted downstream to align with the mod_NNN sequence by accounting for motif insertion sites. Major poly(A) sites have been highlighted by a black vertical line. (**C**) Similar to Figure **B** but with each motif insertion construct plotted separately. Columns designate cumulative plots from different terminator constructs and rows designate cumulative plots from different promoter contexts.

We display the poly(A) sites as the cumulative fraction of poly(A)-site reads mapped at each location downstream of construct stop codons, out of all reads mapped to the terminator (Figure 7B). We confirmed that in other genes the poly(A) site locations were highly reproducible across samples, giving us confidence that changes in reporter poly(A) sites are meaningful (Supplementary Figure S12). Poly(A) sites are in the same relative positions in native loci and the constructs with wild-type terminator (Supplementary Figure S13). Then, we compared 3’end positions of reads between modified and wild-type reporter constructs to determine the poly(A) site usage. In both wild-type and mod_NNN tTSA1 constructs, the same poly(A) sites are used. Surprisingly, tRPS3 mod_NNN constructs appear to be using a novel upstream poly(A) site, that appears in about 50 percent of reads and is located upstream of the 3rd motif insertion site. For tTSA1 constructs the same poly(A) sites are being used in mod_NNN constructs as wildtype (Figure 7B).

Next, we compared changes in poly(A) site usage between constructs with different inserted motifs. We highlight the 2 major poly(A) sites for the tRPS3 constructs and 2 for the tTSA1 constructs (black vertical lines on the mod_NNN constructs in Figure 7B) and track the cumulative fraction of reads upstream of each major site. However, for tRPS3 constructs there is a distinct shift to downstream poly(A) sites in constructs with verified, rather than random, motifs inserted. There are also smaller differences in poly(A) site usage between constructs with different motifs. For tTSA1 constructs there is little change in poly(A) site usage across all constructs.

We then asked whether modifications in the 3’-UTR region impact 5’-3’ degradation following mRNA decapping, using the 5PSeq method targeted to the 3’-end regions of mRNA (49). 5PSeq can detect changing ribosome dynamics through 5’-3’ co-translational degradation, however, the novel modification to 5Pseq here uses an anchored oligo(dT) reverse primer so detects only the poly(A)-site proximal region of the mRNA instead of the entire coding sequence. The 5Pseq counts per gene are highly reproducible between samples (Supplementary Figure S11B), and the abundances of reporter mRNAs from different constructs correlate well between 5PSeq and QuantSeq data (Supplementary Figure S11C). Our 5PSeq data finds no detectable changes in 5’-phosphorylated intermediates between wild-type and modified reporter constructs, and thus does not indicate detectable changes in ribosome dynamics near the 3’ end of transcripts (Supplementary Figure S13, S15). Moreover, the poly(A) site distribution for each construct matches that obtained from QuantSeq data, suggesting that 3’-UTR isoforms are not differentially degraded using this pathway regardless of the motifs inserted. Finally, the abundances of different reporter mRNAs from different constructs correlate well between 5PSeq and QuantSeq data (Supplementary Figures S11C).

Overall, poly(A) site mapping showed that most reporter mRNAs retained the expected poly(A) site and motifs, except for a new alternative poly(A) site in tRPS3 mod_NNN constructs. This highlights the potential for unexpected consequences from composing cis-regulatory elements, even when introducing “random” insertions of no known function.

## DISCUSSION

Our measurements of protein output from promoter-terminator combinations highlight the limitations of assuming that CREs are quantitatively composable. Unsurprisingly, fluorescent protein expression levels are dominated by promoter choice, up to 100 fold, and then terminator choice, up to 5 fold. However, our observations of 1.5 fold change in the relative effect of terminators depending on coding sequence and promoter choice highlight the quantitative limitations of the assumption of composability. Our results echo previous findings of idiosyncratic and context-dependent interactions between CREs (18). Previous work by Kosuri et al. (17) suggests that screening a larger library of CREs would reveal a subset with stronger quantitative interactions.

Terminators taken from co-regulated genes have similar behaviour in our reporter constructs. Terminators that have consistently low expression include tSUN4 and tTOS6, taken from cell wall proteins whose transcripts are bound and regulated by shared RNA-binding proteins (11). Terminators have similar effects when paired with either of the two ribosomal protein promoters pRPS3 or pRPS13, which is consistent with the co-regulation of ribosomal proteins in cells. Further investigations are required into the shared sequence signals and proteins that explain these regulatory patterns and affect the composability of these elements.

We find that the effect of short CREs in 3’UTRs depends on the context of their host construct. The contribution of our selected cis-regulatory motifs to mRNA stability, as inferred from transcriptome-wide analysis, correlates with their effect on the abundance of reporter constructs. However, their contributions to mRNA abundance were inconsistent and dependent on context: TGTAHMNTA has no measurable effect when inserted into tRPS3 but the expected effect in tTSA1, ATATTC has the expected effect in tRPS3 but little effect in tTSA1, and HWNCATTWY can either decrease or increase mRNA levels when removed from tPIR1, depending on the promoter. Also, the two tPIR1 constructs with different mutated HWNCATTWY motifs had different expression levels suggesting HWNCATTWY has a position-dependent effect. The exact sequences of the two HWNCATTWY motif instances did also differ by 4 nucleotides. These disparities in predicted and experimentally verified regulatory behaviour raises questions about current methods for discovering novel cis-regulatory elements and, more generally, for understanding and manipulating regulatory pathways.

The effect of CREs can also depend on environmental context. Our motif search predicted a TGTAAATA motif to be stabilising in the (46) dataset but destabilising in the (38) dataset. TGTAAATA is a known binding motif for Puf3 and carbon deficient conditions trigger Puf3 to degrade its targets (50). Carbon deficient conditions were likely present due to the bespoke CSM-lowURA media used by (38), which is similar to the SC-URA media used in our assays, contrasting with carbon-rich YPD media used by (46). Thus, carbon-dependent regulation by Puf3 could explain the different behaviour of TGTAAATA in the different datasets.

RNA-Seq results confirmed our conclusions on the effect of motif insertions on gene expression. pRPS3-tRPS3 and pTSA1-tTSA1 constructs show similar relative abundances in the qPCR results as the RNA-Seq results. However, pPGK1-tRPS3 results are skewed due to the unexpected low abundance of mod_NNN constructs in the RNA-Seq results, which all other constructs are normalised to. As the QuantSeq protocol we used for our RNA-Seq assay anchors reads to the poly(A) tail of transcripts we were able to determine the effect of inserting motifs on poly(A) site usage. For tTSA1 constructs, inserting the motifs shifted the poly(A) sites 27 nucleotides downstream of their absolute positions in WT, or the total size of the three insertions combined. The sequences and relative usages of the poly(A) sites are not affected by inserting motifs. For tRPS3 constructs, poly(A) sites were again appropriately shifted downstream by the length of the insertions. However, a novel poly(A) site is introduced between the second and third inserted motifs. It is known that efficiency elements in the terminators of yeast genes play an important role in efficient RNA transcription and that cleavage and polyadenylation sites occur a specific distance downstream of the efficiency elements (51). Although we tried to avoid altering efficiency elements in native terminators, the creation of novel poly(A) site is likely due to the inserted motifs extending the distance between the efficiency elements and the native poly(A) sites. Another possible explanation for the novel poly(A) site is that motifs 2 and 3 were inserted into a evolutionarily conserved element of tRps3, as detected by pHastCons (52). This might have disrupted an RBP target site or secondary structure in this region, leading to changes in polyadenylation site choice. Despite this the QuantSeq results do confirm that at least one copy of the motifs of interest are present in every major isoform for all constructs.

### Mechanisms for motif terminator context dependence

This study documents CRE variability with local context, but CRE behavior is also known to vary temporally and spatially in cells, further supporting that CRE functions are not generally decomposable. Our results support long-standing mechanistic observations that CRE contributions depend on the presence of other CREs in the host gene. However, CREs are often regulated by cognate trans-regulatory elements (TREs) that are spatiotemporally regulated themselves. These effects are not separable, because spatiotemporal regulation of gene expression conferred by one CRE can change the interaction of a second CRE with its relevant TREs. For example, a motif may act as the regulatory binding site of a TRE only after the transcript is localised to a time and place where that TRE is present, as we discuss below.

Temporal regulation by successive CREs is common in developmental processes. For example, in *C. elegans* lin-4 is a non-coding RNA gene crucial for regulating cell fates during the early stages of larval development (53). Lin-4 is a small RNA that acts as a TRE by binding to a repeated motif in the 3’UTR of its target mRNA lin-14, and inhibiting the translation of lin-14 (54). Since lin-4 is only expressed at the end of the first larval development stage, the 3’UTR motifs in lin-14 only begin to inhibit its translation late into stage 1, initiating the start of stage 2 (55). The regulation of lin-4 by lin-14 binding to the 3’UTR thus depends on the temporal expression pattern encoded by lin-4’s promoter. Similarly, to establish meiotic chromosome segregation in budding yeast, mRNA encoding cyclin CLB3 is transcribed in stage I of meiosis, but is translationally repressed until stage II of meiosis. CLB3 is translationally repressed via its 5’UTR by the RNA-binding protein Rim4. During the transition to meiosis II, Rim4 is phosphorylated which inhibits binding to CLB3 and enables CLB3 to be translated (56). Here, 5’UTR-specified translational control of CLB3 by Rim4 depends on the timing of promoter-specified transcriptional control.

Spatial regulation by successive CREs is also important, as one CRE can facilitate transcript localisation whilst other CREs and/or TREs confer localisation-dependent regulation. In yeast, the Ash1 protein represses mating-type switching only in daughter cells (57), due to localisation of the ASH1 transcript at the bud tip and subsequent localised translation (58). It is thought that co-transcriptional recruitment of She2 to the ASH1 transcript in the nucleus enables the later recruitment of cytoplasmic factors Khd1/Hek2 and Puf6, factors known for translational repression. Furthermore, successful transport of ASH1 to the bud tip by She2-She3-Myo4 complexes depends on translational repression by Khd1 and Puf6. Later, phosphorylation of Khd1 and Puf6 by budmembrane-localised kinases leads to localised translational activation of the ASH1 mRNP (59, 60).

Another example where the effect of a CRE on a transcript depends on co-localisation with a regulatory kinase comes from the fungal RNA-binding protein Ssd1. Yeast cell wall proteins such as Sun4 and Tos6 are translationally repressed by Ssd1 (61), which binds the 5’UTRs of their transcripts (11, 62). It is thought that these transcripts are translationally activated at bud sites after the phosphorylation of Ssd1 by a localised kinase, Cbk1 (61). There is no evidence that Ssd1 directly acts to transport RNA, so this localised activation presumably depends on localisation that is conferred by other CREs, that recruit other RNA-binding proteins to those transcripts (11), that then in turn recruit transport machinery. We note that post-transcriptional regulation of SUN4 and TOS6 might explain the repressive effect of their terminators when paired with high-expression promoters (Figure 1).

Secondary structures can be integral to motif function, and secondary structure depends on sequences beyond a motif. Again for ASH1, key cis-regulatory elements depend on secondary structure (63). Secondary structure is also essential for Smg recognition elements found in transcript targets of Vts1p (64). In mammalian cells, secondary structures have an overall inhibitory effect on motif functionality (16). In yeast, the presence of stable stem loops within the 3’UTR reduces the binding of known destabilising RBPs (65).

Motif behaviour also depends on CREs across the entire host transcript. Regulation of the ASH1 transcript depends on the concerted action of multiple CREs (66, 67). In other cases, multiple motifs, or multiple copies of the same motif, must occur together to enable recruitment of the effector protein. Recruitment of Puf3 to promote mRNA transcript decay requires two Puf3 motifs in the 3’UTR (68). Indeed, clusters of short degenerate motifs have been proposed as a general regulatory feature of RNA-binding proteins (69).

### Implications for novel cis-regulatory element searches

Motif detection algorithms need to progress beyond the assumption that the presence of a motif alone ensures its functional effect. The effect of a motif can be influenced by factors such as growth conditions and time of measurement. Uncertainty around the active state of a motif could be introduced using latent variables, as used in transcription factor (TF) binding site prediction algorithms to represent the proportion of active TF sites (70,71). Alternatively, non-linear relationships between CREs can be modeled by partition regression techniques (72).

Reductionist search algorithms that attempt to discover the shortest possible sequence contributing to expression may limit the discovery of novel CREs. High resolution maps of protein-RNA interactions are revealing RBPs with gapped, multi-partite motifs (62) or motifs that must be repeated in the same transcript to be effective (68, 73). Common motif searching approaches, such as MEME Suite (14), have extended beyond fixed length, gapless motifs but may still assume motifs occur independently and only once per transcript (74). Although reducing the search constraints to allow longer, gapped motifs could increase the false positive rate, the availability of precise interactome datasets suggests a need for more flexible search algorithms that effectively discover complex motifs.

### Implications for engineering biology

Our results support previous studies showing unpredictability in the expression of combinations of otherwise well-characterised regulatory elements. In the creation of biological circuits, for example, time-consuming directed evolution assays have been implemented to overcome mis-matches in component expression levels (75). The availability of extensively characterised parts libraries now enables large scale searches for CREs with suitable expression. Together with high-throughput assays, pools of combinations of suitable CREs can be quickly screened for the correct expression levels (17).

Our conceptual model of a gene influences the tools we make, and can limit our ability to predict and manipulate gene expression (76). Conceptual models of CREs as independently acting composable parts exemplify these limits. The characterisation of an element’s function in a single context may or may not lead to make generalisable conclusions. Both small- and large-scale studies of CREs should ideally be carried out with multiple combinations of other regulatory elements, in cells in multiple growth conditions, to build a comprehensive understanding of transcriptional and post-transcriptional regulation.

## DATA AVAILABILITY

Plate reader data, RT-qPCR data, and complete analysis code is available at https://github.com/DimmestP/chimera_project_manuscript. RNA-seq data is available on Gene Expression Omnibus GSE200211. Complete code for RNA-seq data analysis is at https://github.com/DimmestP/nextflow_paired_reads_pipeline.

## ACKNOWLEDGMENTS

We thank Naomi Nakayama, Erika Szymanski, Filippo Menolascina, and members of the Wallace lab for discussions on the manuscript. We thank Ivan Clark, Iseabail Farquhar, Luis Montano and Peter Swain for help with the plate reader experiments and analysis. We thank Vicent Pelechano and Yujie Zhang for their generous help teaching us the 5PSeq protocol, obtaining 5PSeq data, and discussing the results. We thank the Genetics Core, Edinburgh Clinical Research Facility, University of Edinburgh for their expertise and assistance obtaining RNA-seq data.

## SUPPLEMENTARY DATA

**Supplementary Table S1.** Variance explained by each type of CRE in the half-life model applied to data from Chan et al (2018). Variance explained was estimated in 3 different ways. The individual variance explained by a linear model containing each type of CRE on its own. The drop in variance explained when one type of CRE is removed from the full model. The cumulative variance explained when each CRE is added to the linear model in sequence; starting with codon usage, then adding motifs and finally 3’UTR length.

| CRE | Individual | Drop | Cumulative |
| --- | --- | --- | --- |
| Codon Usage | 42.0% | -40.5% | 42.0% |
| Motifs | 3.2% | -1.6% | 43.7% |
| 3'UTR Length | 0.6% | 0.0% | 43.7% |

**Supplementary Figure S2.**
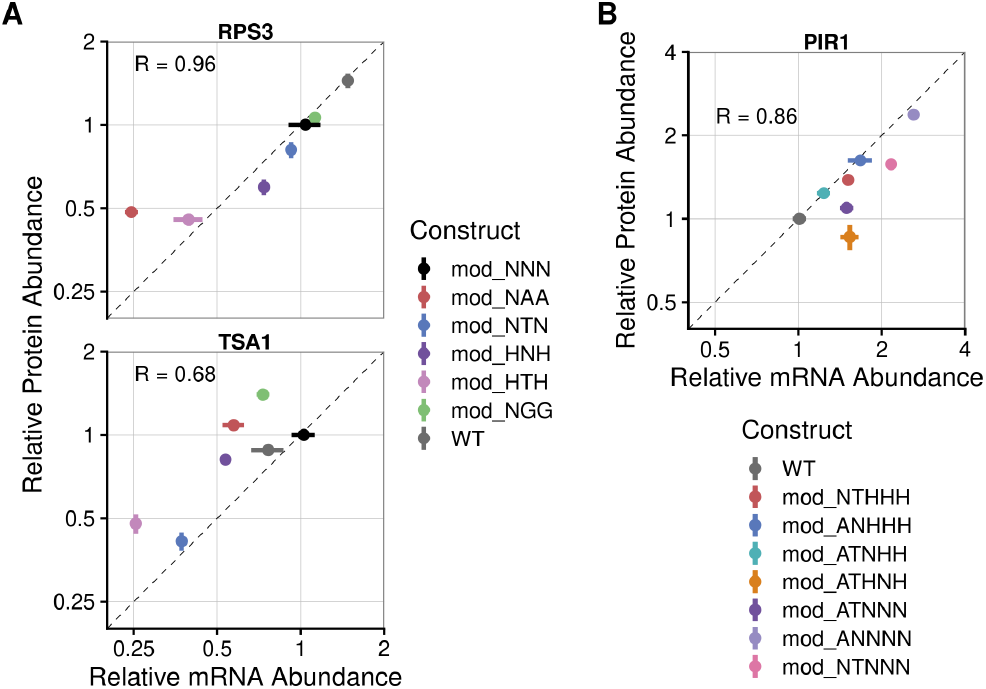
Relative protein abundance correlates with relative mRNA abundance for reporter constructs with modified 3’UTRs. (**A**) Fold changes in RT-qPCR vs fold changes in mCherry fluorescence for all terminator constructs in the pRPS3-mCherry-tRPS3 and pTSA1-mCherry-tTSA1 pairings. Transcript abundance is relative to the mod0 construct of each promoter-terminator pairing. (**B**) Fold changes in RT-qPCR vs fold changes in mCherry fluorescence for all terminator constructs in the pPIR1-mCherry-tPIR1 pairing. Transcript abundance is relative to WT construct of the promoter-terminator pairing. The mean and standard error calculated over 6 biological replicates is plotted for each construct.

**Supplementary Figure S3.**
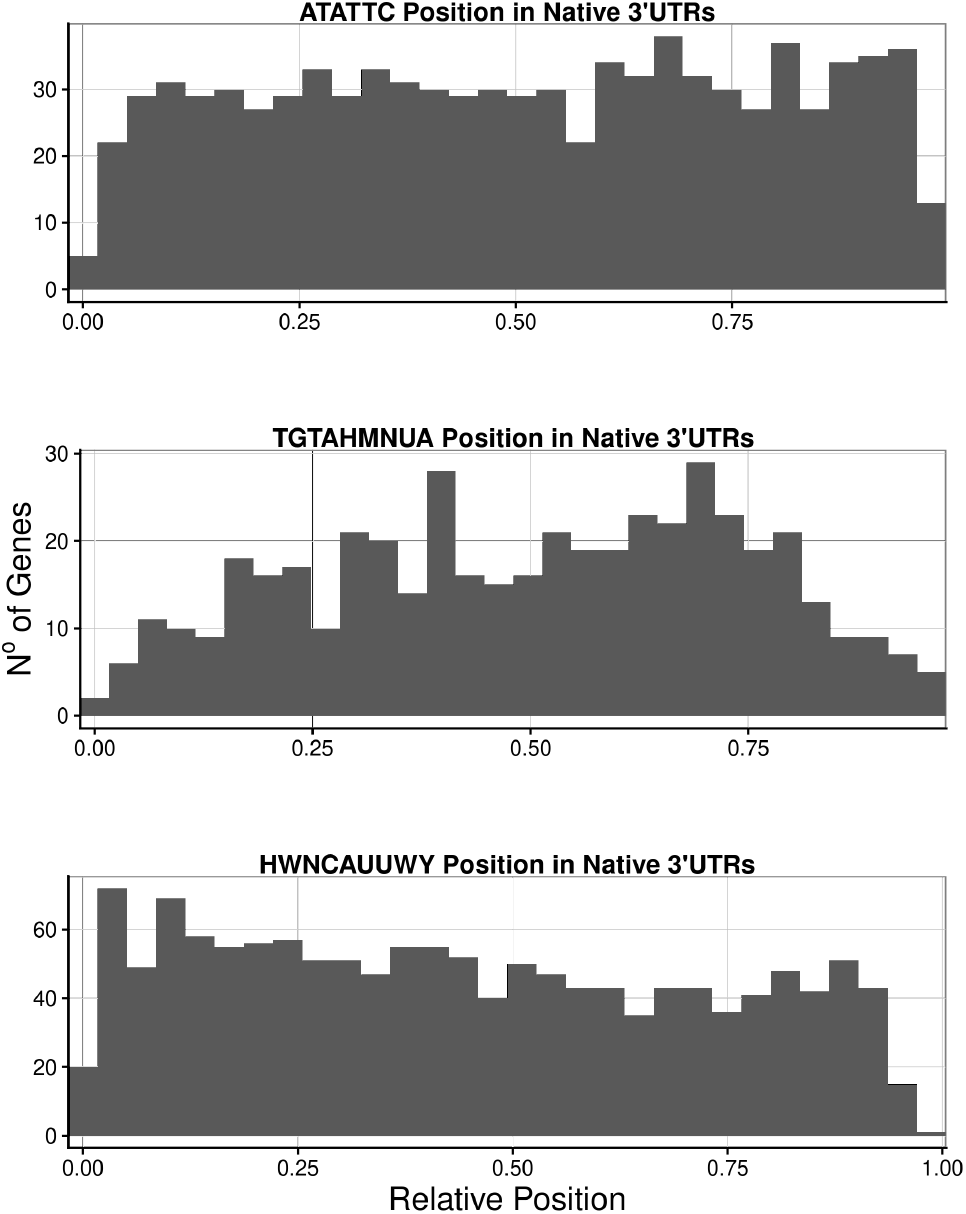
Relative positions of ATATTC, TGTAHMNUA and HWNCAUUWY motifs in native 3’UTRs. Histogram shows the counts of the motif occurrences in 3’UTRs relative to the total length of 3’UTR, where 0 would be starting exactly at the stop codon and 1 would be at the reported poly(A)-site. See (methods or ref to data repository) for details.

**Supplementary Table S4.**
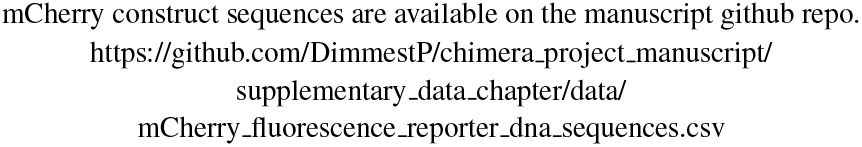
Table showing the DNA sequences for all mCherry reporter constructs.

**Supplementary Table S5.**
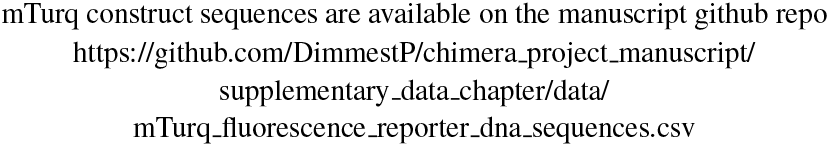
Table showing the DNA sequences for all mTurq reporter constructs.

**Supplementary Table S6.**
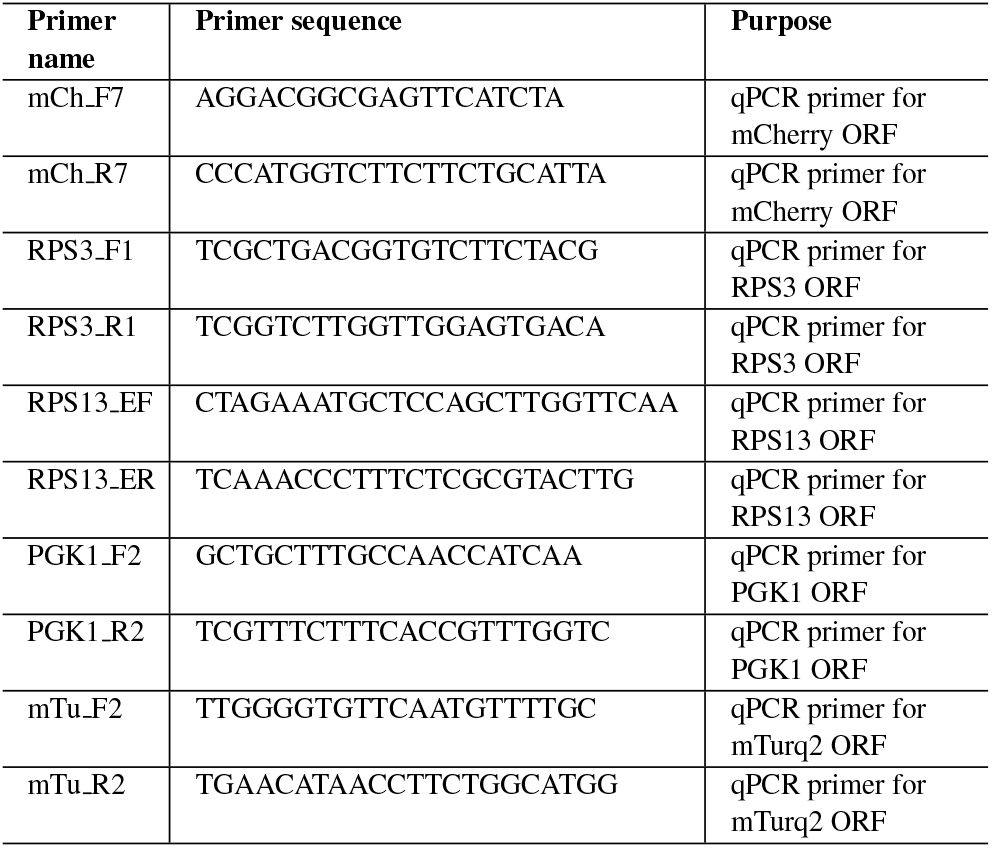
Primer sequences created for all qPCR experiments.

**Supplementary Table S7.**
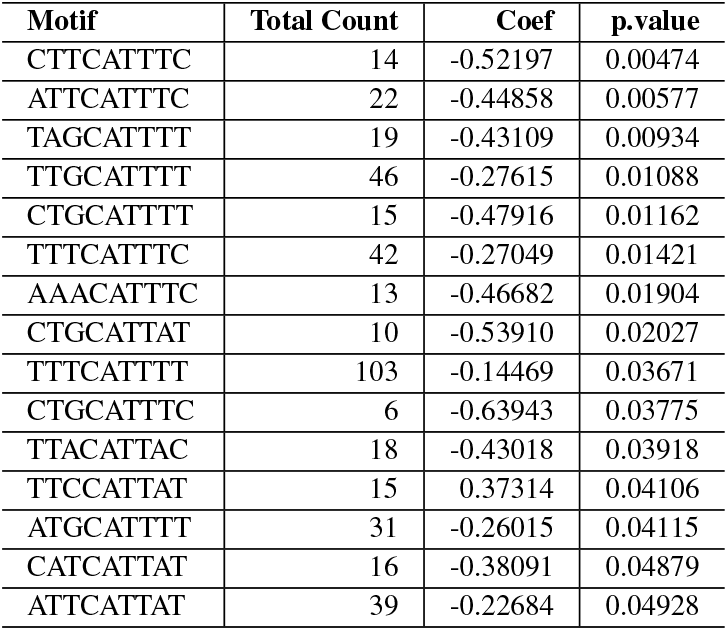
Selecting the HWNCAUUWY motif, TTTCATTTC. Summary of the number of occurences and contributions to a linear model predecting half-life for each possible version of the HWNCAUUWY motif.

**Supplementary Table S8.** Selecting the UGUAHMNUA motif, TGTACAATA. Summary of the number of occurences and contributions to a linear model predecting half-life for each possible version of the UGUAHMNUA motif.

| Motif | Total Count | Coef | p.value |
| --- | --- | --- | --- |
| TGTATATTA | 83 | -0.51920 | 0.00000 |
| TGTATCATA | 17 | -0.67490 | 0.00004 |
| TGTATAATA | 72 | -0.27914 | 0.00066 |
| TGTACCATA | 6 | -0.98705 | 0.00115 |
| TGTACACTA | 16 | -0.56750 | 0.00373 |
| TGTACAATA | 27 | -0.37788 | 0.00551 |
| TGTAACATA | 19 | -0.44503 | 0.00687 |
| TGTATACTA | 36 | -0.30274 | 0.00863 |
| TGTACATTA | 26 | -0.29426 | 0.02791 |

**Supplementary Table S9.** Table showing minimum free energies of 3’UTR constructs with inserted/deleted motifs.

| Terminator | Construct | Minimum free energy (kcal/mol) |
| --- | --- | --- |
| RPS3 | WT | -6.10 |
| RPS3 | mod_NNN | -11.50 |
| RPS3 | mod_NTN | -13.00 |
| RPS3 | mod_NAA | -7.40 |
| RPS3 | mod_NGG | -10.80 |
| RPS3 | mod_HNH | -5.50 |
| RPS3 | mod_HTH | -7.40 |
| TSA1 | WT | -6.90 |
| TSA1 | mod_NNN | -10.14 |
| TSA1 | mod_NTN | -10.60 |
| TSA1 | mod_NAA | -8.90 |
| TSA1 | mod_NGG | -9.80 |
| TSA1 | mod_HNH | -6.10 |
| TSA1 | mod_HTH | -6.67 |
| PIR1 | WT | -28.00 |
| PIR1 | mod_ANHHH | -30.60 |
| PIR1 | mod_NTHHH | -27.70 |
| PIR1 | mod_ATNHH | -28.40 |
| PIR1 | mod_ATHNH | -27.00 |
| PIR1 | mod_ATNNN | -30.40 |
| PIR1 | mod_ANNNN | -32.60 |
| PIR1 | mod_NTNNN | -30.10 |

**Supplementary Figure S10.**
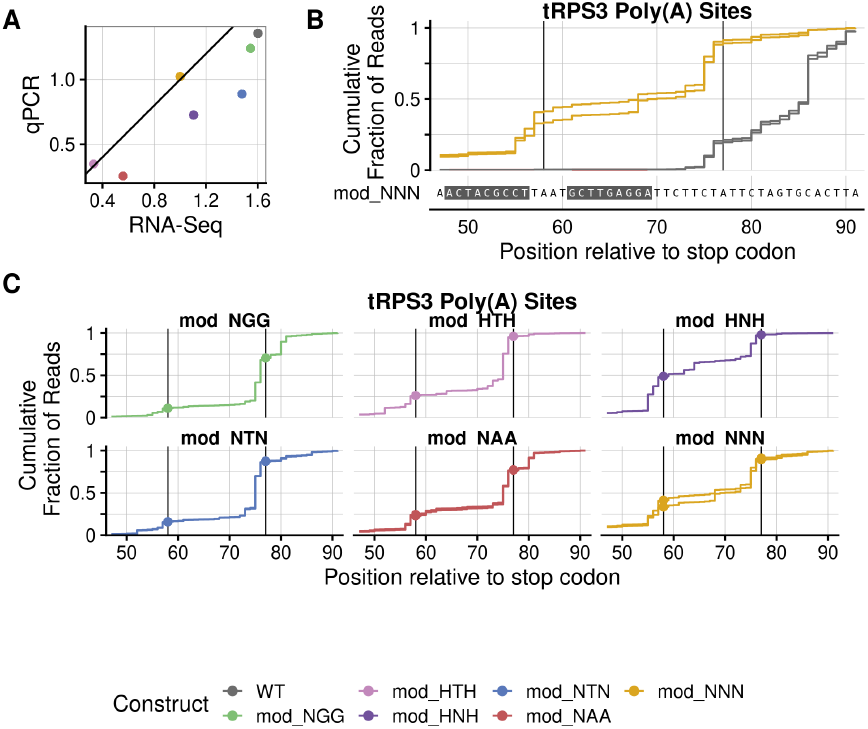
Decapped constructs measured by 5Pseq match mature constructs measured by QuantSeq. pRPS3-tRPS3 constructs were chosen to investigate changes in poly(A) site usage across 5’ decapped transcripts flagged for decay. (**A**) Comparison of construct transcript abundance as independently measured by qPCR and RNA-Seq assays. Transcript abundance was normalised to the median abundance of plasmid URA3, genomic PGK1 and RPS3 transcripts for each construct. Fold change is relative to the mod_NNN construct in each promoter-terminator context. The black diagonal line represents the expected values if RNAseq and qPCR results correlated perfectly. (**B**) Cumulative counts of reads mapped downstream of WT (grey) and mod_NNN (golden) construct stop codons as fraction of total reads mapped to the constructs terminator. WT reads have been shifted downstream to align with the mod_NNN sequence by accounting for motif insertion sites. Major poly(A) sites have been highlighted by a black vertical line. Constructs also used in the QuantSeq analysis have their QuantSeq cumulative graphs plotted in dotted lines. (**C**) Similar to Figure **B** but with each motif insertion construct plotted separately. Columns designate cumulative plots from different terminator constructs.

**Supplementary Figure S11.**
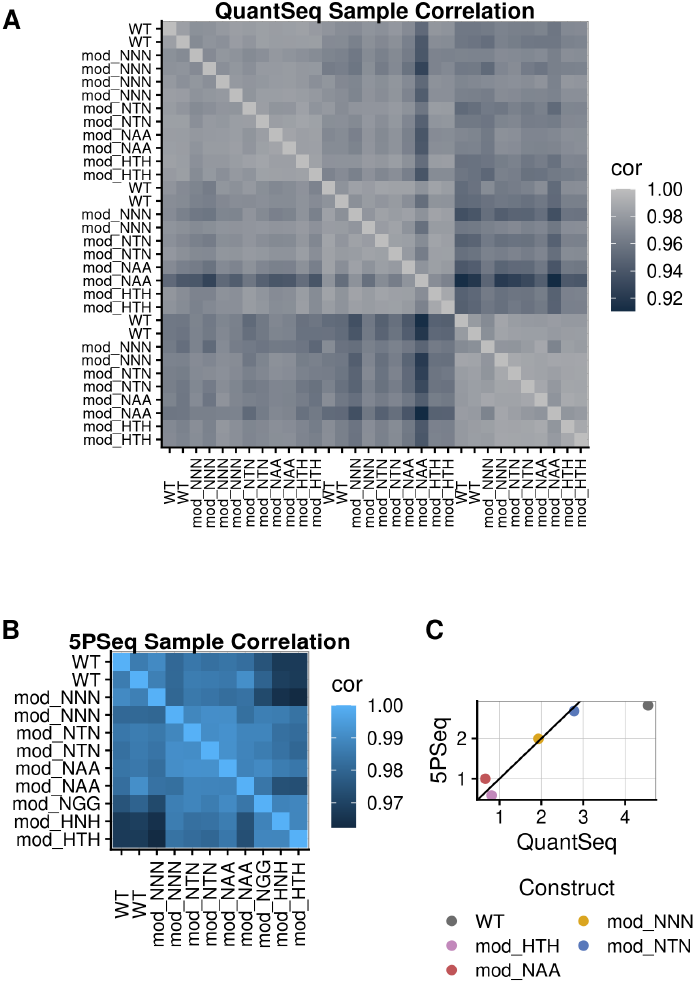
High correlation in transcript counts between samples for both RNA-Seq assays. (**A**) Correlation heat map of DESeq2 normalised log2 pseudocounts across all genes for each sample pair in the QuantSeq assay. From top to bottom on the y axis (left to right on the x-axis) the terminator contexts are pRPS3-tRPS3, pPGK1-tRPS3 and pTSA1-tTSA1. (**B**) Similar correlation heat map for all sample pairs in the 5PSeq assay. (**C**) Comparison of mean log2 abundance of construct mRNA transcripts as measured by 5PSeq and QuantSeq. Only 5 terminator constructs were measured by both methods and only in the pRPS3-tRPS3 context. Transcript abundance for each sample is normalised to the median of the genomic PGK1, TSA1 and RPS3, and the plasmid URA3 genes to match the normalisation used in the qPCR analysis.

**Supplementary Figure S12.**
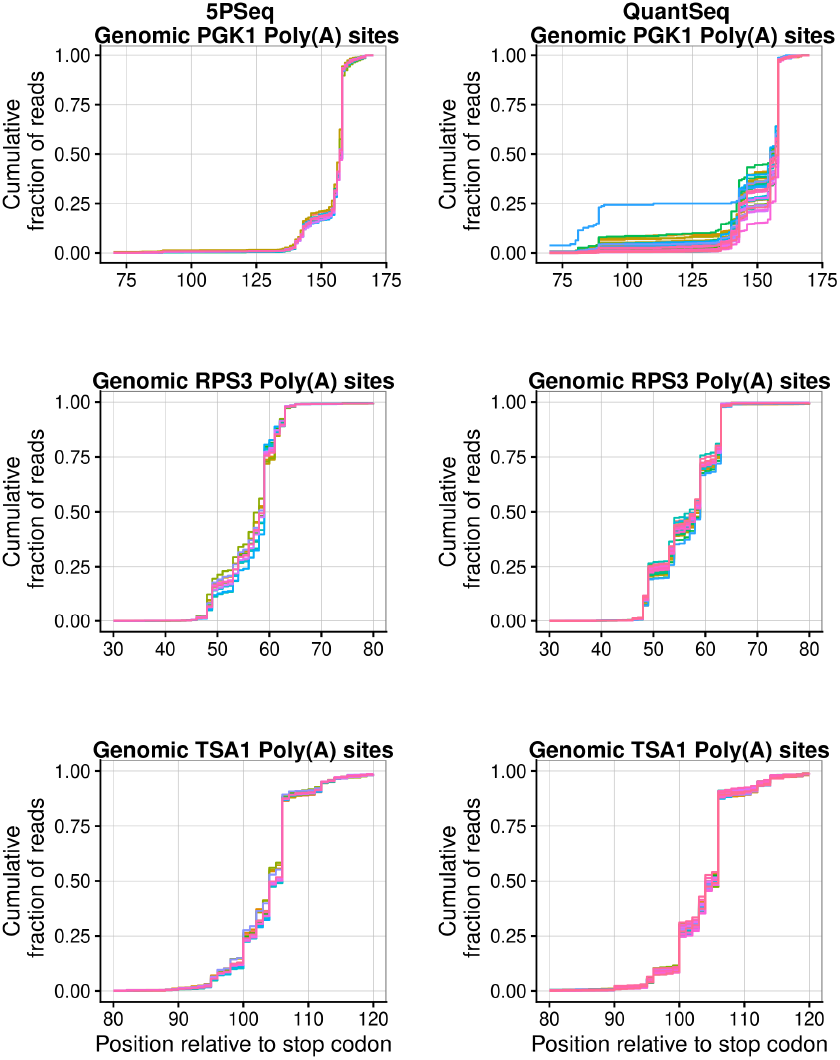
Poly(A) site usage for genomic PGK1, TSA1 and RPS3 terminators remains the same across samples for each RNA-Seq assay. Cumulative counts of reads mapped downstream of native genomic gene stop codons as fraction of total reads mapped to the terminator. Relative usage of poly(A) sites remain similar across samples and across RNA-seq assays. Higher variability in QuantSeq PGK1 poly(A) site usage is due to low overall transcript counts.

**Supplementary Figure S13.**
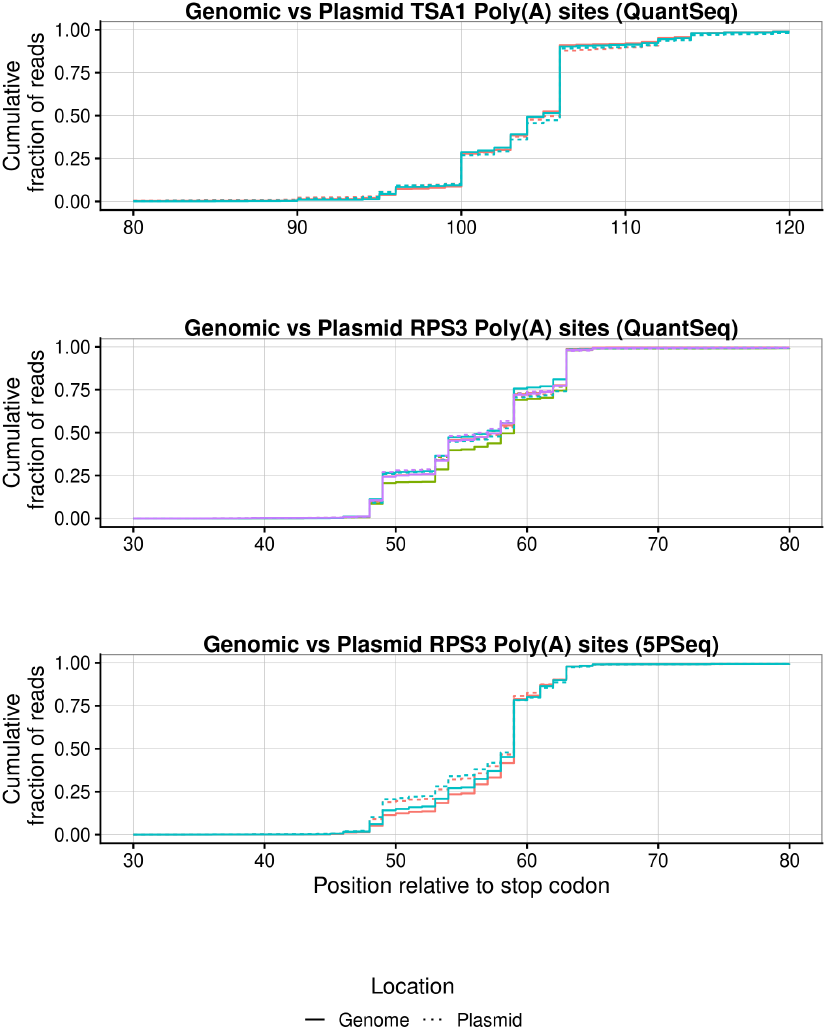
Poly(A) site usage remains the same for genomic TSA1 and RPS3 terminators as for plasmid expressed WT constructs in QuantSeq and 5PSeq. Cumulative counts of reads mapped downstream of WT stop codons for native genomic terminators and plasmid construct terminators as a fraction of total reads mapped to the constructs terminator. Expressing constructs from a low copy number plasmid does not seem to affect poly(A) site usage.

**Supplementary Figure S14.**
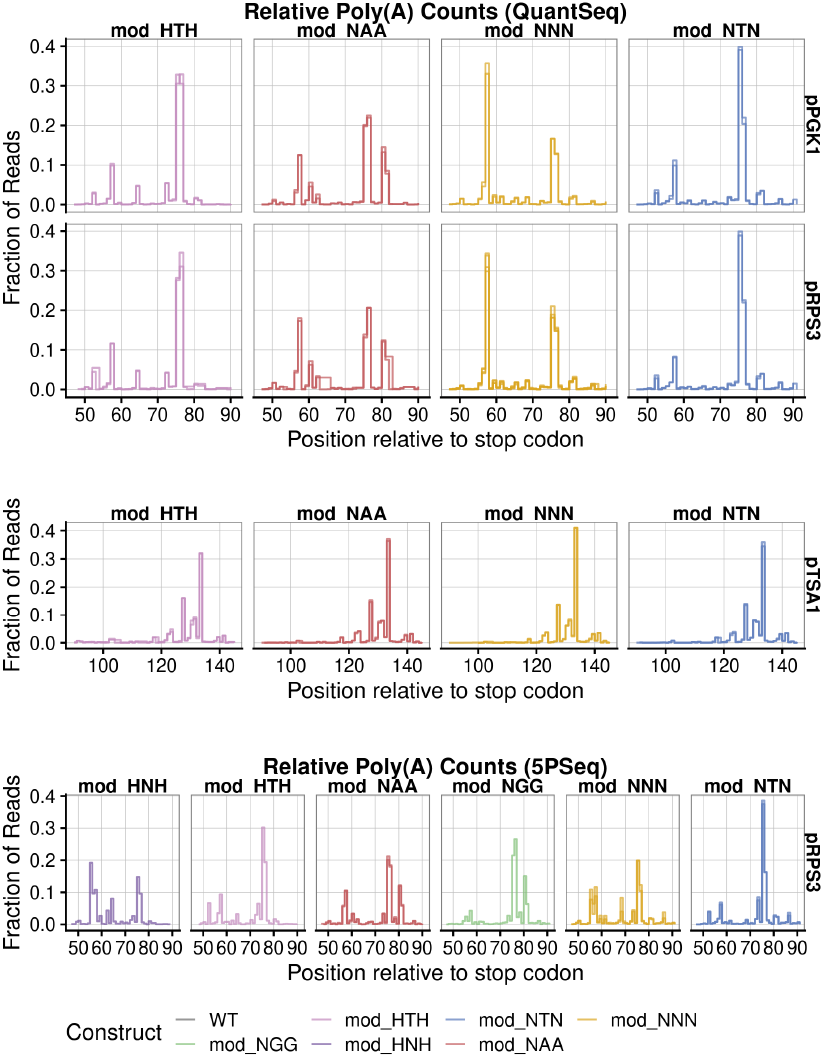
Construct poly(A) site usage across 5PSeq and QuantSeq. Relative counts of reads mapped downstream of construct stop codons as fraction of total reads mapped to the constructs terminator. Peaks represent the position of major poly(A) sites. Replicates are plotted on top of each other where available.

**Supplementary Figure S15.**
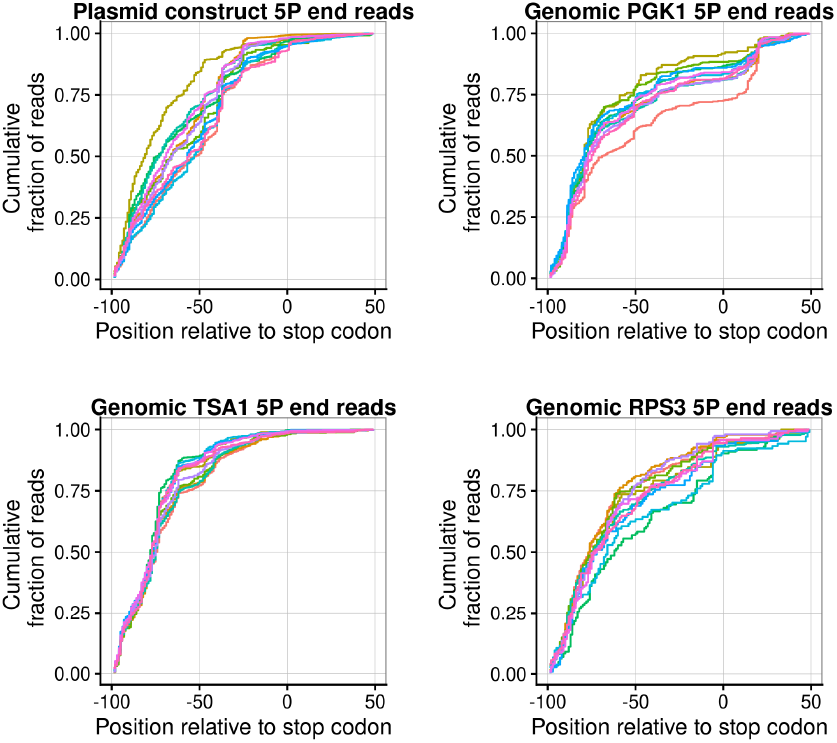
5PSeq data finds no detectable changes in 5’-phosphorylated intermediates between reporter constructs. Relative counts of 5’ end reads of 5’-phosphorylated intermediates as fraction of total 5’ end reads mapped to the terminator. Results for plasmid constructs are plotted alongside selected genomic genes for comparison.

